# Structural Determinants of the Energetic Barrier of Phospholipid Flip-Flop^†^

**DOI:** 10.64898/2026.10.26.761816

**Authors:** Maisa Vuorte, Utkarsh Pandey, Matti Javanainen

**Affiliations:** Computational Physics Laboratory, Tampere University, Korkeakoulunkatu 3, FI-33720 Tampere, Finland

**Author notes:** Supplementary Information (SI) available: mixed-bilayer compositions, simulation protocol, PMF screening, additional results, Bayesian model specification and validation, and description of the data files. See DOI.

## Abstract

Spontaneous phospholipid flip-flop sets the background against which flippases, floppases, and scramblases create and dissipate membrane asymmetry. How the associated free-energy barrier depends on lipid chemistry and on the surrounding membrane remains unclear, because experiments are few and contradictory and previous simulations covered only a small part of the lipidome. Here, we calculate Martini 3 flip-flop free-energy barriers for over 700 membrane–lipid combinations, the largest such dataset to our knowledge. A systematic single component bilayer panel of five headgroups, three linkages, and 35 tail families is complemented by mixed bilayers, from binary mixtures to organelle-inspired compositions, that place the same lipid in different membranes and thus separate the effects of lipid chemistry and the membrane environment. In single component bilayers, the barrier rises with tighter tail packing and depends strongly on headgroup and tail unsaturation but only weakly on linkage. Moving a lipid into a mixed bilayer shifts its barrier as much as most headgroup substitutions do, and the shift follows the change in local tail order. Within a shared membrane, headgroup differences are mostly smaller than between single component bilayers, so part of a headgroup effect arises from the membrane that the lipid forms. A Bayesian model combining chemical descriptors with local membrane structure predicts barriers for withheld lipid families and membranes with cross-validated errors of 2.8 and 3.8 kJ mol^−1^. The calculated trends agree qualitatively with native-lipid vesicle experiments, although Martini 3 underestimates the barrier differences between lipids, gives lower barriers than atomistic models, and overstates the effect of cholesterol.

## 1 Introduction

Phospholipids move between the two leaflets of a membrane by transbilayer translocation, commonly referred to as flip-flop. Spontaneous flip-flop is slow because the polar headgroup must pass through the hydrophobic core of the bilayer. In cells, dedicated lipid-translocating proteins complement this slow process. ^1^ Together, spontaneous and protein-mediated flip-flop determine whether the two leaflets of a membrane differ in composition. The plasma membrane of mammalian cells is asymmetric, with phosphatidylserine (PS) and phosphatidylethanolamine (PE) concentrated in the cytoplasmic leaflet and phosphatidylcholine (PC) and sphingomyelin enriched in the exoplasmic leaflet, and the two leaflets also differ in lipid chain unsaturation and in their structural properties. ^2^ This asymmetry is maintained by ATP-dependent flippases and floppases and dissipated by scramblases, which are activated during platelet activation and apoptosis. ^1,3,4^

In the endoplasmic reticulum (ER), by contrast, lipids are constitutively scrambled between the leaflets without energy input. ^1^ All of these processes act against the background of spontaneous flip-flop, which determines how quickly an asymmetric distribution relaxes without protein assistance.

Measured rates of spontaneous flip-flop depend more on the experimental setup than on the membrane composition. In asymmetric DPPC vesicles, fluid-phase half-times ranged from a day at 65 °C to multiple days at 50 °C, whereas no exchange was detected in the gel phase within a span of ten days. ^5^ In contrast, measurements on planar supported bilayers ^6^ give rates that are orders of magnitude faster, which has been attributed to defects caused by the support. ^5^ Labelled lipids can also distort both the flip-flop rate and the associated activation energetics. ^7^ For example, a spin-labelled PC flipped about ten times more slowly than native DPPC, ^6^ whereas an NBD-labelled PC flipped faster than the native lipids around it. ^8^ Native-lipid methods avoid this problem but typically require isotopic contrast, as in time-resolved neutron scattering. ^9^ Simulations under well-controlled conditions can therefore help to resolve the trends between lipids and to separate them from the effects of the experimental technique and model system. ^10,11^

Still, a handful of systematic experiments show that both the chemistry of the lipid and its membrane environment matter. Pyrene-labelled phosphatidylglycerol (PG), phosphatidic acid (PA), PE, and PC probes in POPC vesicles crossed at rates increasing in the order PC *<* PG *<* PA *<* PE, with PE translocation being at least an order of magnitude faster than that of PC. ^12^ In supported bilayers of native lipids, the activation free energy of flip-flop increases with chain length and with the packing and compression modulus of the membrane. ^6,7,13,14^ In vesicles of native lipids at 37 °C, DMPC and POPA flipped with half-times of about 6 and 7 h, whereas POPC flip-flop was not detected, implying a half-time of more than 1000 h. ^15^ In the same study, 20 and 40 mol% cholesterol increased the DMPC half-time to 22 h and more than 100 h, respectively.

Computational studies reveal how the surrounding membrane takes part in the crossing. In atomistic simulations, simply forcing a DPPC headgroup into the bilayer interior nucleates a transmembrane water pore. ^16^ Once a pore has formed, lipids can cross the membrane within tens of nanoseconds with their head-group hydrated, which suggests that pore formation is the rate-limiting step. ^17,18^ United-atom simulations at 323 K gave barriers that range from 16 kJ mol^−1^ for dilauroyl-PC (DLPC) to about 90 kJ mol^−1^ for POPC and DOPC. Water pores formed in the saturated PC bilayers but not in POPC or DOPC, and the barriers mainly reflected the cost of deforming the bilayer. ^19^ A recent transition-path sampling study identified four distinct crossing mechanisms, namely water nanopores, water threads across locally thinned membranes, water nanodroplets, and dry tunnelling, depending on the lipid and the model. ^20^ Importantly, cellular membranes differ widely in properties that are likely to affect these pathways. One key difference is the leaflet asymmetry of the plasma membrane compared with the nearly symmetric ER membrane. Moreover, cholesterol makes up only about 5% of ER lipids but 30–40% of plasma-membrane lipids, which strongly affects lipid packing and lateral heterogeneity. ^21^ In united-atom DPPC bilayers, 20 and 40 mol% cholesterol raised the flip-flop barrier from 80 to 106 and 111 kJ mol^−1^, respectively. ^22^ Both the chemistry of the flip-flopping lipid and the membrane environment are therefore likely to determine spontaneous flip-flop rates.

Coarse-grained models enable much larger chemical surveys of the lipidome. Martini 3 maps about four heavy atoms onto a single bead, ^23^ and its refined lipidome improves agreement with measured areas per lipid, bilayer thicknesses, main transition temperatures, and cholesterol-dependent phase behaviour. ^24^ A recent study compared Martini 3 and atomistic CHARMM36m barriers for six lipids with different headgroups in a POPC membrane and for five single component PC bilayers. ^25,26^ The Martini 3 barriers were 20–70% lower for most lipids, and they hardly changed between saturated PC bilayers with 12–18 carbons per chain. ^25,26^ In some cases, the flip-flop process also differs qualitatively between Martini 3 and atomistic models, as DMPC crosses the bilayer without a water pore only in Martini 3. ^20^ Martini 3 thus appears suited to studying trends within the model rather than to providing quantitative estimates of experimental barriers or rates.

The large lipidome and efficiency of Martini 3 make it possible to probe how the barrier depends on the chemistry of the flip-flopping lipid and on the structural properties of the membrane. Martini 3 reproduces membrane structure well, ^24^ whereas its coarse-grained description of individual lipid chemistries may be less accurate. High-throughput Martini 3 data could also train machine learning models that predict barriers beyond the training set, for example for membranes whose composition changes with disease, ^27^ diet, ^28^ or external conditions. ^29^ However, to our knowledge, the largest previous comparison of flip-flop barriers covered only eleven lipids. ^25^

Here, we extend this scope by calculating Martini 3 flip-flop free-energy barriers for a systematic panel of over 600 single component bilayers covering five glycerophospholipid headgroups, three linkages, and 35 tail families. We then examine how the membrane environment modifies the barrier using (1) binary and ternary mixtures, (2) cholesterol series that range from disordered to highly ordered packing, and (3) compositions inspired by cellular membranes, including asymmetric bilayers. Combining these datasets, we compare the effect of changing the membrane environment with that of changing the flip-flopping lipid. Finally, we test whether lipid chemistry and membrane structure together can predict the barriers for lipids and membranes beyond the training set.

## 2 Computational methods

### 2.1 Coarse-grained simulations of lipid bilayers

We calculated the structural properties of each bilayer with unbiased simulations and its flip-flop barriers with biased simulations.

Our systematic single component bilayer panel consisted of two-tail glycerophospholipids with PA, PC, PE, PG, or PS headgroups, ester, ether, or plasmalogen linkages, and acyl chain lengths spanning 8–22 carbons with zero to six double bonds per chain. This set was supplemented by a small number of phosphatidylinositol (PI), sphingomyelin (SM), and bis(monoacylglycero)phosphate (BMP) bilayers. Lipids with the same numbers of carbons and double bonds in their acyl chains form *a tail family*. The mixed bilayers comprised binary and ternary phospholipid mixtures, binary DMPC/cholesterol and POPC/cholesterol series, and compositions inspired by the plasma membrane, ER, Golgi apparatus, lysosome, endosome, mitochondrial inner and outer membranes, and a bacterial inner membrane. The plasma-membrane and mitochondrial compositions include asymmetric bilayers together with leaflet-symmetric controls, in which both leaflets have the composition of one leaflet of the asymmetric bilayer. Fig. 1 summarizes the chemical diversity of the simulated lipids and the simulation approach, and shows examples of a single component and a mixed bilayer. The ER, Golgi, lysosome, endosome, and plasma-membrane compositions were adapted from the all-atom organelle models of Xu *et al*., ^30^ the asymmetric mitochondrial compositions from the Martini 3 models of Brown *et al*., ^31^ the simplified mitochondrial inner-membrane composition from Makowski *et al*., ^32^ and the bacterial inner-membrane composition from the *E. coli* model of Corey et al. ^33^ These are compositional representations rather than models of complete organelle membranes. The leaflet compositions are listed in Table S1 of the Supplementary Information (SI).

**Fig 1.**
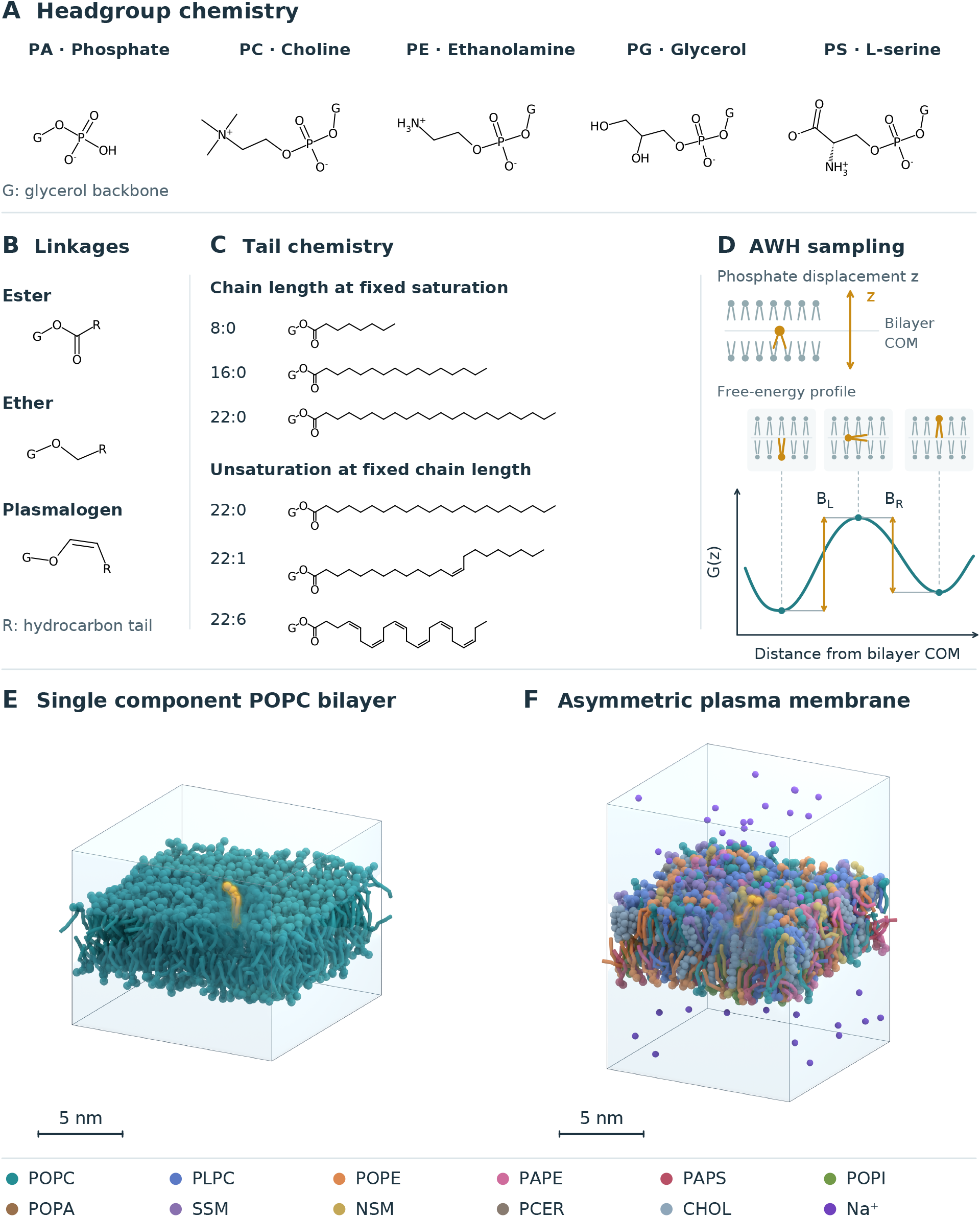
Chemical variation of the simulated lipids and simulation approach. (A–C) Chemical variables of the dataset. (A) Headgroups of phosphatidic acid (PA), phosphatidylcholine (PC), phosphatidylethanolamine (PE), phosphatidylglycerol (PG), and phosphatidylserine (PS), with G denoting the glycerol backbone. (B) Ester, ether, and plasmalogen (vinyl ether) linkages, with R denoting the hydrocarbon tail. (C) Examples of tails that differ in length at fixed saturation and in the number of *cis* double bonds at fixed length, labelled by the numbers of carbon atoms and double bonds per chain. The dataset also contains phosphatidylinositol (PI), sphingomyelin (SM), and bis(monoacylglycero)phosphate (BMP) lipids. (D) AWH sampling along the displacement *z* of the phosphate bead from the centre of mass of the bilayer, and a schematic free-energy profile with the side barriers *B*_L_ and *B*_R_ from the two leaflet minima to the central maximum. (E, F) Snapshots of the single component POPC bilayer and the asymmetric plasma-membrane composition, with the flip-flopping lipid shown in orange. Water is shown as a translucent blue volume.

All simulations were carried out with GROMACS (version 2025.4) ^34^ using the Martini 3 force field ^23^ with the version 3.2 lipid parameters of the refined Martini 3 lipidome. ^24^ Bilayers were built with insane, ^35^ extended with custom definitions for Martini 3 lipids that the builder did not support at the time, and solvated with standard Martini water. Single component bilayers contained 300 lipids per leaflet, whereas mixed bilayers were built with lateral dimensions of 14 × 14 nm, so that their numbers of molecules depend on the composition and, in asymmetric bilayers, can differ between the two leaflets (Table S1). Na^+^ or Cl^−^ counterions were added only to neutralize the net charge of the membrane. Single component and mixed bilayers were simulated with the same protocol and settings (Table S2). They were energy-minimized in three steepest-descent stages and then pre-equilibrated for 35 ns in six stages, with the time step increasing from 2 to 20 fs, using a protocol modified from that of CHARMM-GUI Martini Maker. ^36^ Production simulations used a 20 fs time step, periodic boundary conditions, and Verlet neighbour lists. ^37^ Electrostatic interactions were treated with the reaction-field method, with a relative dielectric constant of 15 and a cutoff of 1.1 nm. ^38^ Lennard-Jones interactions were cut off at 1.1 nm with the potential shifted to zero at the cutoff distance. The temperature was maintained at 310.15 K with the stochastic velocity-rescaling thermostat (coupling time 1 ps), ^39^ coupling the membrane and the solvent separately. The pressure was coupled semi-isotropically to 1 bar (compressibility 3 ×10^−4^ bar^−1^), using the Berendsen barostat ^40^ in the first five pre-equilibration stages, the Parrinello–Rahman barostat ^41^ in the last, and the stochastic cell-rescaling barostat (coupling time 4 ps) ^42^ in the production simulations. The unbiased and biased simulations used the same settings and were started independently from the pre-equilibrated bilayers, with new velocities generated at 310.15 K. All bilayers were simulated without bias for 10 µs, and structural properties were averaged over the final 5 µs, with standard errors from ten 0.5 µs blocks.

### 2.2 Estimation of flip-flop free energies

Flip-flop was sampled with the accelerated weight histogram (AWH) method, ^43,44^ which adaptively builds up a bias along a chosen reaction coordinate until a target distribution, here a uniform one, is sampled. The converged bias provides an estimate of the potential of mean force (PMF) along the reaction coordinate. Here, the coordinate was the displacement of the phosphate bead of the flip-flopping lipid along the bilayer normal, relative to the centre of mass of the bilayer. The lipids crossed the membrane 514±228 (SD) times on average during the 15 µs-long AWH runs, which largely eliminates hysteresis. The number of crossings varied widely between lipids, which contributes to the differences in PMF convergence between them.

The convolved form of the AWH bias was used, with a harmonic coupling force constant of 10^4^ kJ mol^−1^ nm^−2^, a uniform target distribution, an initial error estimate of 10 kJ mol^−1^, and a diffusion estimate of 10^−3^ nm^2^ ps^−1^. The sampled interval was symmetric with respect to the bilayer centre and was set for each bilayer based on its thickness, so that the free-energy minima in the headgroup regions of both leaflets were fully covered (SI Section S1.2). Each PMF was obtained from a single AWH simulation in which bias was applied to a single lipid. For each PMF, Δ*G*(*z*), the side barriers *B*_L_ and *B*_R_ are the heights of its maximum above the minima in the left (*z <* 0) and right (*z >* 0) leaflets (Fig. 1D). Their mean, *B*_ave_ = (*B*_L_ + *B*_R_)*/*2, is the mean flip-flop barrier.

In a symmetric bilayer, *B*_L_ and *B*_R_ provide two estimates for the barrier, whereas in an asymmetric bilayer they differ by the free-energy difference between the two leaflet basins, *B*_L_ − *B*_R_ = Δ*G*(*z*_R_) − Δ*G*(*z*_L_). Averaging the two side barriers removes this leaflet preference, so *B*_ave_ measures the cost of reaching the transition region irrespective of direction. The directional barriers in the asymmetric bilayers are listed in Table S6.

PMF profiles were accepted based on temporal convergence and symmetry criteria (Fig. S1 and SI Section S1.3). Between cumulative snapshots separated by at least 1 µs, neither the barrier height nor the root mean square of the profile was allowed to change by more than 2 kJ mol^−1^. In the single component bilayers, the phospholipid and cholesterol mixtures, the leaflet-symmetric controls, and the simplified mitochondrial inner-membrane composition, the two side barriers were also required to agree within 10 kJ mol^−1^. Profiles that met these criteria passed the PMF stability screen. The accepted side barriers differed by a median of 0.7 kJ mol^−1^ in single component and 2.3 kJ mol^−1^ in mixed bilayers.

### 2.3 Structural descriptors and identification of gel-like bilayers

Area per lipid, bilayer thickness, and tail order were obtained from the unbiased simulations. For a tail bond vector at an angle *θ* to the bilayer normal (*z* axis), the orientational order parameter is

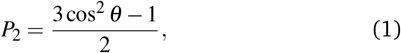

where 0 corresponds to random orientations and 1 to perfect alignment with the membrane normal. We used two complementary definitions for *P*_2_. Bulk tail order averages *P*_2_ over all consecutive tail-bead vectors, whereas interfacial tail order uses only the vector from the first to the second tail bead after the linker, *i*.*e*. it reports only the orientation of the uppermost tail segment shared by all studied lipid species.

For single component bilayers, area per lipid is the box area *L*_*x*_*L*_*y*_ divided by the number of lipids *N* per leaflet, whereas thickness is calculated from the inter-leaflet distance of the average phosphate bead positions along the membrane normal (*z* axis). For mixed bilayers, the descriptors were averaged over the in-plane neighbourhood within 2 nm of each molecule of the flip-flopping species. Area was obtained from a leaflet-wise Voronoi tessellation and thickness from the local headgroup positions. These definitions capture the environment sampled by a particular lipid molecule, so they can differ for two flip-flopping lipid species in the same membrane.

Gel-like bilayers were identified from their structure alone and excluded from all analyses. Area, thickness, and both order parameters of the 608 single component bilayers with complete structural data were standardized, and their first principal component, which explained 89.25% of the variance and weights the four descriptors almost equally, served as a packing coordinate. A two-component Gaussian mixture fitted along this coordinate classified 123 bilayers as gel-like, with a probability weight of at least 0.5 for the ordered component, and the remaining 485 as fluid-like (Fig. S2). This classification is based on structure only and is not a thermodynamic phase assignment, and borderline systems would require diffusion, positional order, or temperature scans to be classified accurately. Projected onto the same coordinate, seven combinations in four mixed bilayers, namely DMPC with 30 and 40 mol% cholesterol, POPC with 30 mol% cholesterol, and the outer-leaflet control of the plasma-membrane composition, also fall into the gel-like region. These systems were retained, because high tail order in cholesterol-containing membranes indicates a liquid-ordered rather than a gel phase.

### 2.4 Statistical analysis and predictive surrogate model

To test whether chemistry and structure predict barriers outside the data used for fitting, a Bayesian regression model was fitted to the logarithm of the barrier,

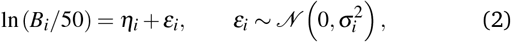

where *B*_*i*_ is in kJ mol^−1^. Due to the logarithmic scale, inputs act multiplicatively on the barrier itself. The mean *η*_*i*_ contains 10 inputs, namely the headgroup and linkage categories, five numerical tail descriptors (total tail carbons, total double bonds, doublebond asymmetry between the two chains, unsaturation position, and mapped tail bending), and three structural descriptors from the unbiased simulations (bilayer thickness, bulk tail order, and interfacial tail order). Limited nonlinearity is allowed through spline terms in total tail carbons, thickness, and interfacial order, and through interactions of the headgroup with interfacial order and of total tail carbons with thickness. Each tail family and each mixed bilayer receives its own offset, and the mixed bilayers also have a common mean offset, corrections to the structural slopes, and their own residual standard deviation *σ*_*i*_. No coefficient is fitted for an individual lipid. All coefficients have normal priors centred on zero, which shrink weakly supported terms, and the posterior and the 95% predictive intervals are obtained analytically. Area per lipid was also examined as an input but was discarded, because it is defined differently for single component and mixed bilayers and added no resolved predictive information (SI Section S3.5). The full model specification is given in SI Section S3.

Predictive performance was assessed by grouped cross-validation, in which related observations are withheld together. ^45^ In the family test, all headgroup and linkage variants of a tail family were withheld together (five folds over 35 families). In the membrane test, each mixed bilayer was withheld together with any leaflet-symmetric controls derived from it (five folds over 25 groups), while all single component data remained in training.

This test assesses predictions for known lipids in a new membrane environment. Accuracy is reported as the root mean square error (RMSE) of the predicted median, with 95% bootstrap intervals over withheld groups, and as the coverage of the 95% predictive intervals (Table S10). The information carried by each input variable was assessed by removing it together with its spline, interaction, and mixed-slope terms, refitting the model, and recording the change in RMSE (Fig. 5C).

## 3 Results and discussion

### 3.1 Dataset and sampling domain

We first discarded gel-like bilayers and those with poor PMF stability. Of the 608 single component bilayers, 429 passed both screens. Of the 152 mixed-bilayer profiles, which cover 138 membrane–lipid combinations, 80 passed the PMF stability screen. After the barriers of profiles started from opposite leaflets were averaged, these profiles provided 74 membrane–lipid combinations in 29 host bilayers (Table S3). Fig. 2A–C shows the composition of the dataset by headgroup, tail saturation, and linkage. The accepted single component bilayers cover all five headgroups and three linkages in 35 tail families. In contrast, the mixed bilayers cover a much narrower chemistry, with 18 PA, PC, PE, PG, and PS lipids and eight PI, SM, and BMP lipids. PC is the flip-flopping lipid in 38 of the 74 combinations (Fig. 2A and Table S1), and the mixed bilayers contain no ether or plasmalogen lipids (Fig. 2C).

**Fig 2.**
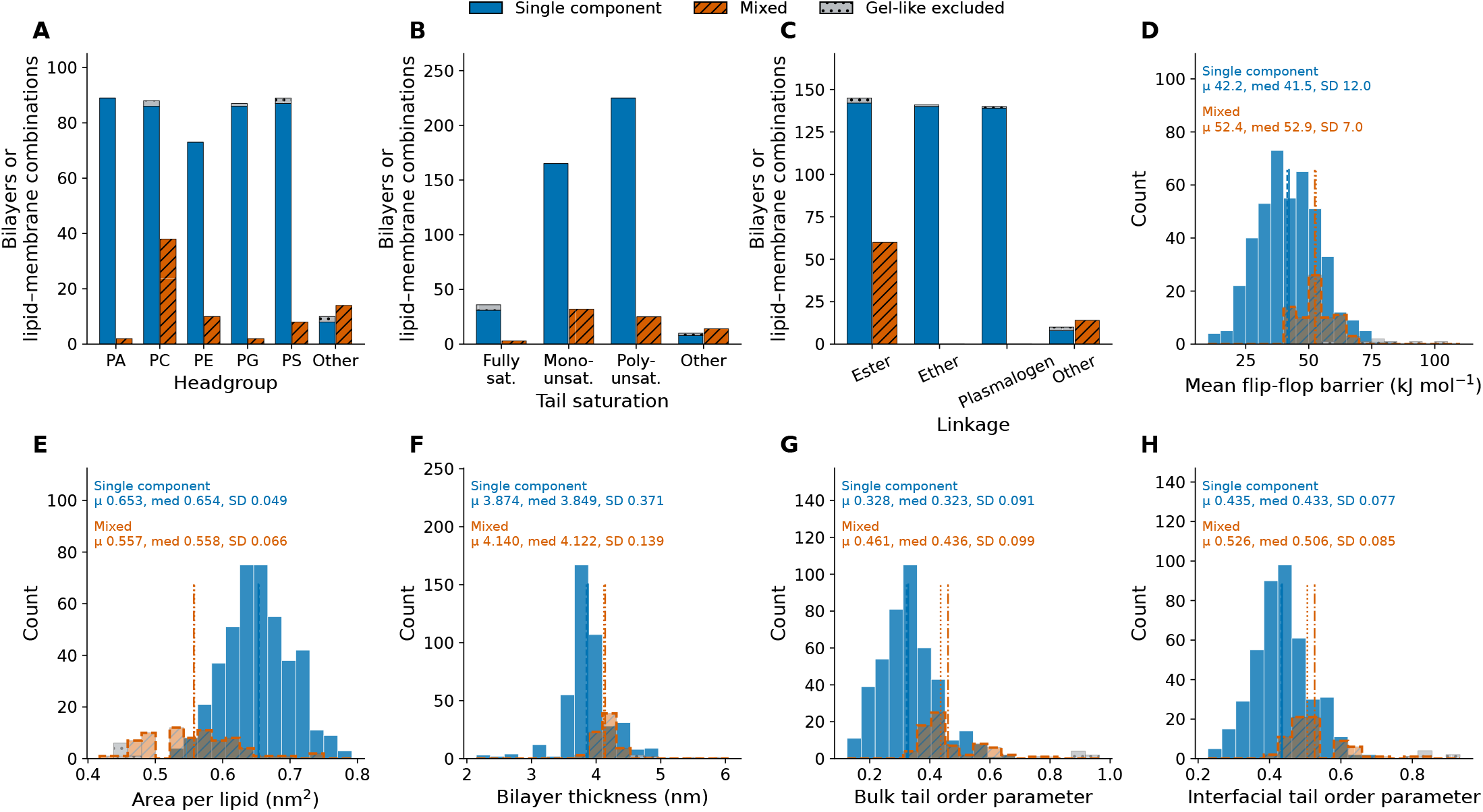
Composition and structural properties of the screened dataset. (A–C) Numbers of single component bilayers (blue) and mixed membrane– lipid combinations (orange, hatched) by headgroup, tail saturation, and linkage; “Other” comprises PI, SM, and BMP lipids. (D–H) Distributions of the mean flip-flop barrier, *B*_ave_, area per lipid, bilayer thickness, bulk tail order, and interfacial tail order for 429 single component bilayers and 74 mixed combinations, with means (*µ*), medians (med), and sample standard deviations (SD). Vertical lines mark the means (solid, single component; dash-dotted, mixed) and medians (dashed and dotted, respectively). Grey dotted bars show the seven gel-like single component bilayers whose profiles passed the PMF stability screen; these are excluded from all analyses and summary statistics.

As expected, gel-like single component bilayers had much higher apparent barriers than fluid-like ones (mean of 117 against 42 kJ mol^−1^), consistent with the absence of detectable gel-phase exchange in DPPC vesicles. ^5^ The associated PMF profiles, however, mostly did not stabilize within the sampled time, and only 7 of the 123 gel-like bilayers passed the PMF stability screen (Table S3). Among fluid-like single component bilayers, 56 of 485 were excluded, more than half of them with short saturated tails of 8–12 carbons per chain. The nine accepted bilayers with 8:0 or 10:0 chains are the thinnest bilayers in the dataset (2.27–

2.68 nm; Fig. 3B). Under experimental conditions, such short-chain lipids may form micelles, and the unsaturated PE lipids may form inverted hexagonal phases. ^46^ Here, their barriers refer only to the planar bilayers imposed by the periodic simulation box. Our absolute barriers differ from earlier calculations with the same model. The barriers of the single component POPC and DMPC bilayers, 53.7 and 47.4 kJ mol^−1^, are lower than the 59 and 62 kJ mol^−1^ that Kroutil *et al*. obtained with umbrella sampling. ^25^ Both studies use the phosphate position along the bilayer normal as the reaction coordinate, so the difference likely reflects the salt concentration (0.15 M NaCl against counterions only), the version of the lipid parameters, and the definition used for barrier extraction. In addition, umbrella sampling windows seeded from a single pulling trajectory can retain hysteresis in slow orthogonal degrees of freedom, ^47^ whereas AWH samples these repeatedly as the lipid crosses the membrane. Absolute barriers therefore depend on the simulation protocol.

**Fig 3.**
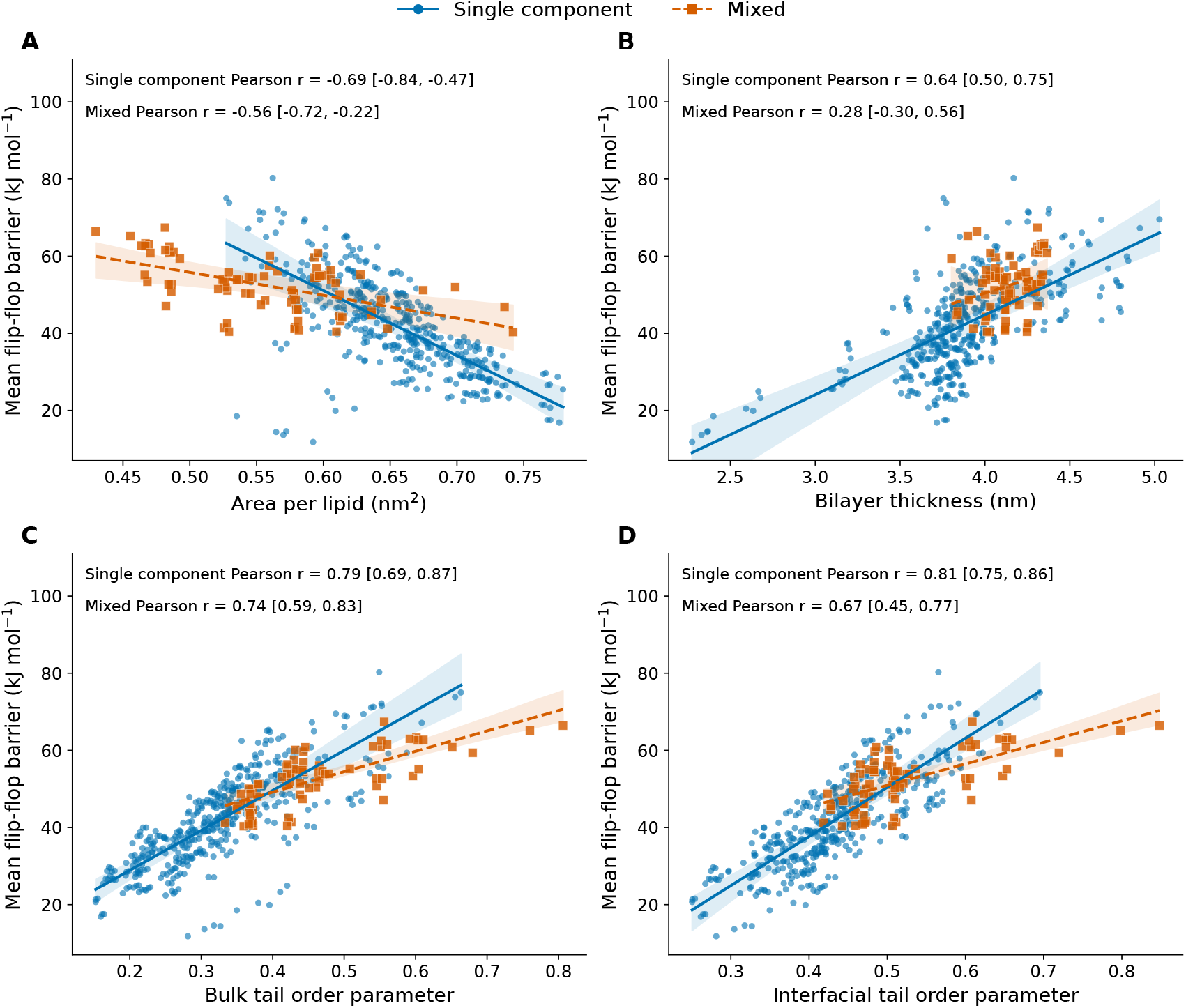
Association of the mean flip-flop barrier, *B*_ave_, with (A) area per lipid, (B) bilayer thickness, (C) bulk tail order, and (D) interfacial tail order. Circles and solid lines show 429 single component bilayers, and squares and dashed lines the 74 mixed-bilayer combinations. Lines are ordinary least-squares fits, and shaded bands are 95% bootstrap intervals for the fitted mean. Pearson coefficients, *r*, are given with 95% cluster-bootstrap intervals over tail families and over mixed bilayers grouped with their leaflet-symmetric controls.

The mixed bilayers span a smaller range of barriers and thick-nesses than the single component bilayers (Fig. 2D and F), and their areas per lipid are not directly comparable, because the two populations use different definitions (Fig. 2E). The mixed bilayers also reach higher tail order than any fluid-like single component bilayer, namely in the DMPC mixtures with 20–40 mol% cholesterol (Fig. 2G and H).

### 3.2 Packing and chemistry trends in single component bilayers

Across single component bilayers, the flip-flop barrier increases with tail order and thickness and decreases with area per lipid (Fig. 3). The two order parameters show the strongest associations (Fig. 3C and D), followed by area per lipid (Fig. 3A). Most bilayers (89%) are 3.5–4.5 nm thick, and within this range the dataset-wide fit describes the trend poorly (Fig. 3B). The association with interfacial order holds within each headgroup, with *r* between 0.78 (PS) and 0.90 (PE). However, the four structural descriptors are strongly correlated with one another (SI Section S2.1), so the strengths of their associations do not rank independent physical effects.

A tighter, more ordered bilayer is expected to resist the local deformation needed to accommodate a polar headgroup in its interior. Earlier atomistic and united-atom studies attributed large differences between PC barriers to such deformation ^19^ and found that DSPC crosses through a locally thinned membrane. ^20^ In line with this, the activation free energy of flip-flop in supported bilayers increases with lipid packing ^13^ and with the compression modulus of the membrane. ^7,14^ The mean descriptors used here cannot, however, separate a deformation cost from headgroup dehydration or defect formation. In Martini 3, DMPC crosses without the water nanopores that mediate crossing in atomistic simulations. ^20^ This may be because pore formation is more costly in Martini than in atomistic models, ^18,24^ so Martini 3 may understate the role of hydrated defects.

Fig. 4A compares the headgroups of lipids with identical tails. PC serves as the reference, because it has the most tail compositions in the panel, is the most common flip-flopping lipid in the mixed bilayers, and is the most abundant phospholipid class in most of the eukaryotic compositions considered here (Table S1). At matched tails and linkage, PE has the highest barriers, 12.7 kJ mol^−1^ above PC in the median, and its barrier is higher in all 65 matched pairs (Table S4). This trend, along with PS being intermediate between PC and PE, was also observed in our recent work on lipid scramblases, where the host membrane for each flip-flopping lipid was POPC. ^48^ These results agree qualitatively with the slower flip-flop of DSPE than of DSPC in gel-phase single component bilayers, ^13^ although the bilayers compared here are structurally fluid-like. Pyrene-labelled PE probes in POPC vesicles, in contrast, crossed faster than PC probes, ^12^ an order that Martini 3 does not reproduce even in mixed bilayers, where

**Fig 4.**
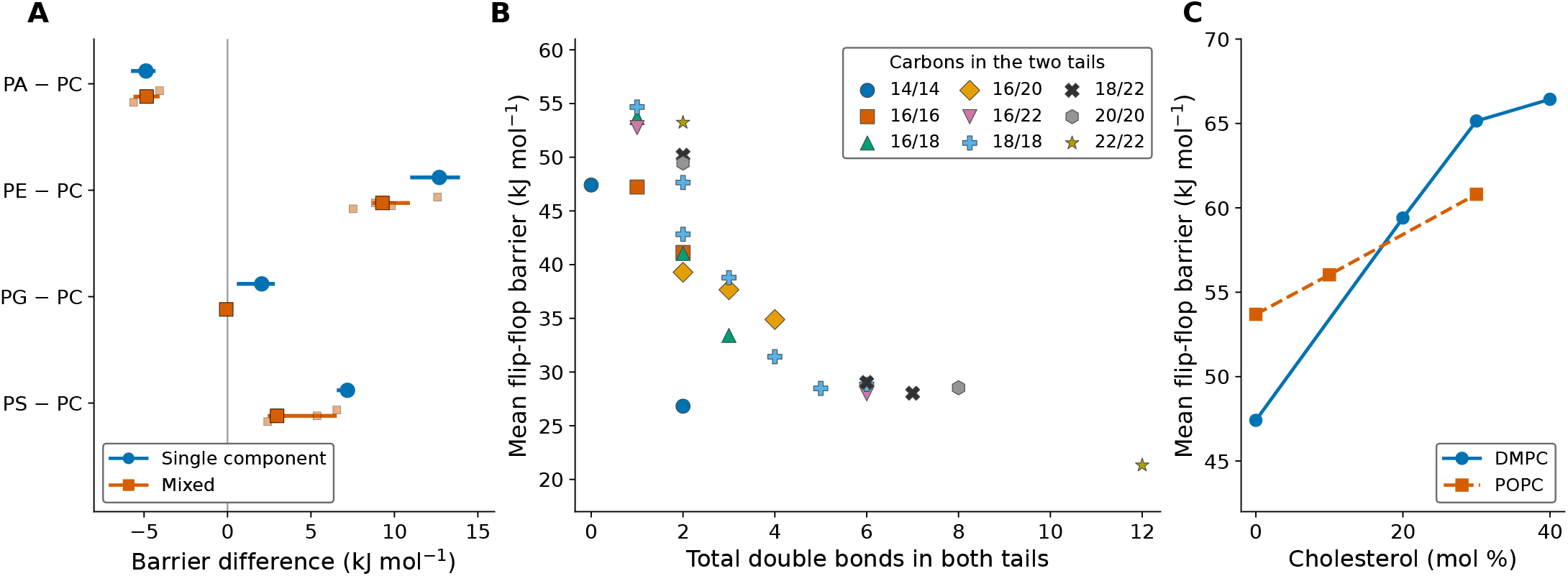
Chemical and membrane effects on the mean flip-flop barrier, *B*_ave_. (A) Headgroup effect relative to PC, *i*.*e*. the barrier difference between a lipid and its PC analogue with the same tails and linkage. Circles compare the two lipids in separate single component bilayers, and squares compare them within the same mixed bilayer. Large symbols show medians with 95% cluster-bootstrap intervals, and small squares show the individual mixed-bilayer pairs. (B) Barriers of single component ester-PC bilayers against the total number of double bonds, for nine series with fixed carbon numbers in the two tails. The two 18/18 points with two double bonds (blue crosses) are di-18:1-PC (upper) and 18:0/18:2-PC (lower), and the 18:1(11-*cis*) isomer is not shown. (C) Barriers of DMPC and POPC in bilayers with cholesterol.

**Fig 5.**
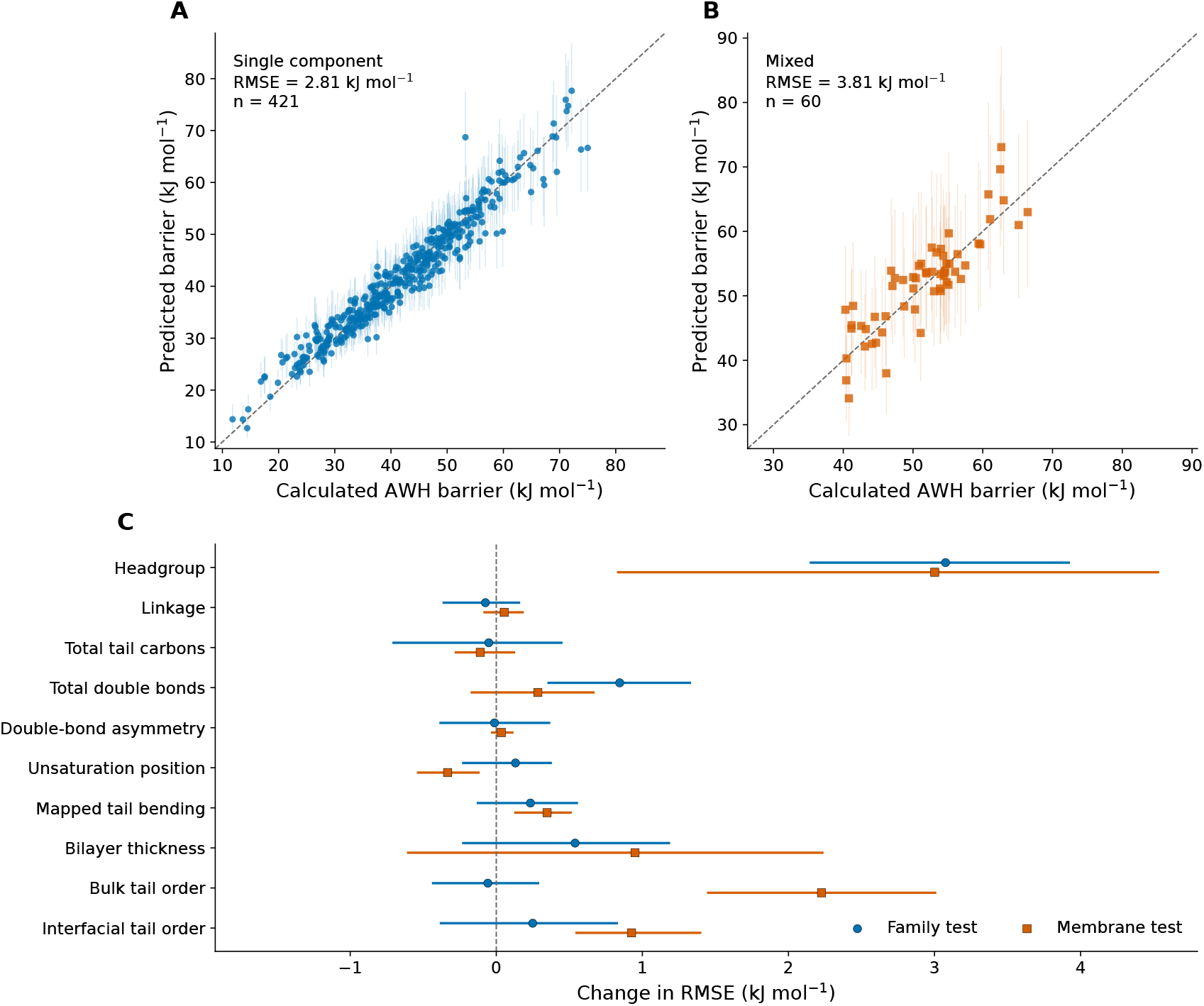
Prediction of mean flip-flop barriers by the Bayesian model. (A) Single component bilayers with their tail family withheld (*n* = 421) and (B) mixed-bilayer combinations with their bilayer withheld (*n* = 60), shown with 95% predictive intervals; dashed lines mark equality. (C) Change in RMSE when each input is removed and the model refitted, for the family test (circles) and the membrane test (squares), with 95% paired bootstrap intervals; positive values indicate poorer predictions.

PE barriers remain above the PC ones (Fig. 4A). Labelled probes can, however, perturb flip-flop energetics, and their barriers need not follow those of the surrounding lipids. ^7^ Atomistic PE bilayers form hydrogen-bond networks between headgroups, ^49^ which Martini represents only through effective Lennard-Jones interactions, so the PE–PC contrast in Martini 3 cannot be attributed specifically to hydrogen bonding.

The anionic headgroups do not follow a common trend. Relative to PC, the median barrier is 7.2 kJ mol^−1^ higher for PS and 2.1 kJ mol^−1^ higher for PG, but 4.9 kJ mol^−1^ lower for PA (Fig. 4A), so formal charge alone does not determine the order of the barriers. PA has a smaller headgroup, and its lower barrier agrees qualitatively with native-lipid experiments, in which POPA vesicles showed flip-flop with a half-time of about 7 h, whereas no flip-flop was detected in POPC vesicles. ^15^ To compare our barriers with measured rates, we assume that two lipids cross with equal prefactors in the Kramers rate theory, ^50^ so that a ratio of half-times corresponds to a barrier difference of *B*_1_ − *B*_2_ = *RT* ln(*t*_1*/*2,1_*/t*_1*/*2,2_), *i*.*e*. 5.9 kJ mol^−1^ for a ten-fold ratio at 310.15 K. This assumption is untested, because the prefactor depends on the diffusivity along the coordinate and on recrossing of the transition region, ^50,51^ and it is unlikely to hold for mechanistically different pathways, *e*.*g*., those involving co-permeating or pore-forming water molecules. With this assumption, the measured half-times imply a barrier difference of more than 12 kJ mol^−1^, compared with 7.0 kJ mol^−1^ between the single component POPA and POPC barriers obtained here (46.7 against 53.7 kJ mol^−1^; Table S7). The Martini topologies have fixed charge states, whereas PA translocation depends on protonation among other contributing effects. ^52^ PG translocation correlates with the density of PG headgroups and the ionic strength, ^53^ whereas PS translocation in supported bilayers is slowed about five-fold by the addition of Ca^2+. 54^ The results for anionic lipids are therefore limited to the modelled charge states and to counterion-only conditions simulated here.

Linkage has a smaller effect than most headgroup substitutions. Relative to ester linkages, ether linkages lower the barrier by a median of 1.9 kJ mol^−1^ at matched headgroup and tails, with lower barriers in 127 of the 129 pairs (Table S4), whereas plasmalogen linkages raise it by only 0.3 kJ mol^−1^ (95% CI 0.2–0.4), a resolved difference that is smaller than the median difference between the two side barriers of an individual profile. To our knowledge, no atomistic or experimental barriers are available to test whether Martini 3 captures the effect of the linkage.

Tail chemistry has a larger effect on the barrier than linkage. At fixed per-chain carbon counts, additional double bonds lower the barrier in 530 of 541 matched comparisons (median −11.1 kJ mol^−1^; Table S4). Longer chains in turn raise the barrier. Extending both chains of a symmetric monounsaturated lipid by two carbons raises the barrier in 74 of 78 matched comparisons (median +3.9 kJ mol^−1^). Notably, this increase levels off with chain length, from a median of 12.4 kJ mol^−1^ between di-14:1 and di-16:1 lipids to 3.2 kJ mol^−1^ between di-20:1 and di-22:1 lipids, as also seen for the ester PCs in Fig. 4B. Saturated PCs, of which only DMPC is shown in Fig. 4B, behave similarly. Replacing one 14:0 chain of DMPC with a 16:0 chain raises the barrier from 47.4 to 57.4–59.4 kJ mol^−1^, depending on the chain position, whereas extending this chain further to 18:0 has practically no effect (57.9–59.2 kJ mol^−1^). The position of the double bond also matters. Moving the *cis* double bond of 18:1 chains from position 9 to position 11 lowers the barrier in all 15 matched headgroup–linkage pairs, by 1.7–6.6 kJ mol^−1^, although interfacial tail order increases, opposite to the dataset-wide trend in Fig. 3D. A single order parameter therefore cannot replace chemical lipid descriptors, which motivates combining the two in the predictive surrogate model.

These chain-length effects differ from earlier Martini 3 calculations. With refined Martini 3 parameters, Kroutil *et al*. obtained essentially the same barrier for di-12:0-PC (DLPC), di-14:0-PC (DMPC), di-16:0-PC (DPPC), and di-18:0-PC (DSPC), as well as for DOPC. ^25,26^ The two studies differ most for the thin DMPC bilayer, for which their barrier is 14.6 kJ mol^−1^ higher than ours. For POPC, the difference is only 5.3 kJ mol^−1^, and our barriers of 57.4–59.4 kJ mol^−1^ for the 14:0/16:0 and 14:0/18:0 PCs are close to the 59–62 kJ mol^−1^ they obtained for POPC and DMPC. Thin bilayers deform strongly as the headgroup crosses, ^19^ so hysteresis may affect umbrella sampling and AWH simulations differently and lead to different converged barriers. In addition, DPPC and DSPC bilayers are expected to be gel-like at 310 K, as their main transition temperatures are 41 and 55 °C, ^55,56^ and DPPC is indeed gel-like in our simulations.

In united-atom simulations at 323 K, the barrier rises steeply with chain length, from 16 kJ mol^−1^ for di-12:0-PC to 40 kJ mol^−1^ for DMPC and 80 kJ mol^−1^ for DPPC, ^19^ and in gel-phase supported bilayers, the activation free energy of saturated PCs increases by about 14 kJ mol^−1^ per two methylene groups per chain. ^7^ In contrast, two additional carbons in the *sn*-1 chain of pyrene-labelled PCs in POPC vesicles only doubled the flip-flop half-time, which corresponds to about 1.8 kJ mol^−1. 12^ Much of the chain-length effect in single component bilayers may therefore arise from the membrane rather than from the flip-flopping lipid itself. However, Martini 3 compresses differences that combine chain length and unsaturation. The DMPC and DOPC barriers are almost equal here (47.4 and 47.6 kJ mol^−1^) and differ by only 5 kJ mol^−1^ in the calculations of Kroutil *et al*. (62 and 57 kJ mol^−1^), ^25^ whereas the united-atom barriers are 40 and 94 kJ mol^−1. 19^ Similarly, native DMPC vesicles showed flip-flop with a half-time of 350 min at 37 °C, whereas none was detected in POPC vesicles, ^15^ which implies a barrier difference of more than 13 kJ mol^−1^ with equal prefactors, against 6.3 kJ mol^−1^ here. The experimental effect of unsaturation is itself unclear, as polyunsaturated PC bilayers are thinner and have a lower bending rigidity, ^57^ which would be expected to favour flip-flop, whereas unsaturated lipids in the opposing leaflet suppressed DPPC flip-flop in supported bilayers. ^58^ Whether unsaturation lowers the barrier as strongly in experiments therefore remains open.

### 3.3 Separating effects of lipid chemistry from the membrane environment

In a single component bilayer, changing the flip-flopping lipid also changes the surrounding membrane. Mixed bilayers instead place the same lipid in different membrane environments (Fig. S4). Moving a lipid from its single component bilayer into a mixed one shifted its barrier by a median of 4.2 kJ mol^−1^ in either direction, which is of the same order as the median effects of the PA, PG, and PS headgroups relative to PC (Fig. 4A). The largest shifts were −13.7 kJ mol^−1^ for SAPI in the lysosome-inspired bilayer and +19.0 kJ mol^−1^ for DMPC with 40 mol% cholesterol. The shifts follow the accompanying change in bulk tail order (*r* = 0.83, 95% CI 0.77–0.90) and, more weakly, those in interfacial order, area, and thickness (Fig. S3). The dependence on tail order is not caused by cholesterol alone, as the correlation remains for the 18 combinations in cholesterol-free mixed bilayers (*r* = 0.89, 95% CI 0.63–0.97). The correlation also remains when the single component references are replaced by each lipid’s mean barrier across the mixed bilayers (SI Section S2.1). The barrier of a lipid thus depends on the packing of its surroundings, and its single component barrier is a poor estimate of its barrier in a mixture. Across mixed bilayers, moreover, the barrier rises only about half as steeply with tail order as across single component bilayers and shows no clear dependence on thickness (Fig. 3), so single component relationships do not transfer directly to mixtures.

The membrane also affects headgroup contrasts. In 13 comparisons, a PA, PE, PG, or PS lipid and its PC analogue with the same tails and linkage share a mixed bilayer (Fig. 4A and Table S5). These contrasts keep the signs found in single component bilayers, apart from the near-zero POPG–POPC contrast, but are smaller in magnitude in 10 of the 13 cases. For example, POPE exceeds POPC by 17.6 kJ mol^−1^ in single component bilayers but by only 7.5–9.8 kJ mol^−1^ in the two plasma-membrane compositions. The latter is in line with the difference of 11 kJ mol^−1^ calculated for these lipids in a POPC host membrane in two recent studies. ^25,48^ Barriers obtained by Kroutil *et al*. for single lipids in POPC bilayers agree in sign with the same-membrane contrasts, including a slightly lower barrier for POPG than for POPC. ^25^ Part of a single component headgroup contrast therefore appears to arise from the packing of the membrane that the lipid forms rather than from the headgroup itself. Experimentally, the large difference between DSPE and DSPC flip-flop rates in single component gel-phase bilayers disappears in mixed DSPE/DSPC bilayers, ^13^ and the DPPS barrier increases with the PS content of DSPC bilayers. ^7^ The comparisons are, however, few and mostly in PC-rich membranes, as more than half of the mixed-bilayer PE profiles failed the PMF stability screen (SI Section S1.3).

Cholesterol raises the barriers of DMPC and POPC strongly and monotonically with its concentration (Fig. 4C). United-atom DPPC simulations also showed that cholesterol raises the barrier and suppresses pore-mediated crossing. ^22^ However, in native-lipid DMPC vesicles at 37 °C, the temperature simulated here, the measured half-times imply a barrier increase of only 3.4 kJ mol^−1^ at 20 mol% cholesterol with equal prefactors, ^15^ compared with 12.0 kJ mol^−1^ in Martini 3 (Table S7), which suggests that Martini 3 might overstate the effect of cholesterol.

Polyunsaturated lipids act in the opposite direction. In 50:50 mixtures with PLPC and SAPC, the POPC barrier drops from 53.7 to 45.6 and 41.2 kJ mol^−1^, as the bulk tail order around POPC decreases from 0.40 to 0.35 and 0.33, respectively. These shifts are close to those expected from the change in tail order alone (Fig. S3), whereas the thickness around POPC barely changes in the SAPC mixture. The organelle-inspired compositions combine both effects. The POPC, PLPC, and POPA barriers are 6.6– 10.6 kJ mol^−1^ higher in the plasma-membrane compositions with 29–35 mol% of cholesterol than in the ER-inspired composition with only 5%. This difference, which likely arises from the ordering effect of cholesterol, is comparable to the headgroup contrasts in Fig. 4A.

In asymmetric bilayers, the numbers of lipids that insane places in the two leaflets of the fixed box area can create differential stress, which alters leaflet packing ^59^ and is not necessarily relieved by the redistribution of cholesterol. ^60^ Excluding the three asymmetric compositions, however, leaves the correlation between barrier shifts and bulk tail order essentially unchanged (*r*= 0.82), and the asymmetric barriers mostly lie within 3 kJ mol^−1^ of those in the leaflet-symmetric controls (Table S6). Their side barriers also contain the leaflet preference of the flip-flopping lipid, which *B*_ave_ averages out (SI Section S2.3).

### 3.4 Prediction of flip-flop from the surrogate model

The Bayesian model predicted the barriers of withheld tail families of single component bilayers with an RMSE of 2.8 kJ mol^−1^ (95% CI 2.2–3.4) and those of withheld mixed bilayers with an RMSE of 3.8 kJ mol^−1^ (2.8–4.4; Fig. 5A and B). These errors are small compared with the spread of the calculated flip-flop barriers, which in the model dataset range from 11.8 to 75.0 kJ mol^−1^ in single component bilayers and from 40.3 to 66.4 kJ mol^−1^ in mixed-bilayer combinations (standard deviations of 11.8 and 6.7 kJ mol^−1^). The model thus captures most of the variation in single component bilayers (*R*^2^ = 0.94) and two-thirds of that in mixed bilayers (*R*^2^ = 0.67; Table S10). It estimates barriers of new lipid families reliably and those of new membranes more coarsely. Its 95% predictive intervals contain 96.7% of the mixed-bilayer combinations and 94.6% of the single component bilayers other than PA, but only 80.9% of the PA lipids (Fig. S5). Mixed-bilayer training data substantially improve predictions for new membranes. Trained only on single component bilayers, the regression model predicted the 60 mixed-bilayer combinations with PA, PC, PE, PG, or PS headgroups with an RMSE of 5.9 kJ mol^−1^, compared with 3.8 kJ mol^−1^ when the other mixed bilayers were included in training (Table S11). Relationships learned from single component bilayers therefore do not transfer directly to mixtures, consistent with the shallower slopes of the mixed bilayers in Fig. 3.

Removing an input and refitting shows how much of its information the other model inputs cannot replace (Fig. 5C). The headgroup is the most important input, and removing it raises the error by approximately 3 kJ mol^−1^ for both withheld tail families and withheld mixed bilayers. The headgroup thus carries information beyond that of the structural inputs. This is also in line with the large headgroup effects at matched tails (Fig. 4A). The structural inputs matter mainly for new membranes. For withheld mixed bilayers, removing all three raises the RMSE by 3.9 kJ mol^−1^ (SI Section S3.5), mostly through bulk tail order, whereas bulk tail order is dispensable for withheld tail families. In a single component bilayer, tail order follows from the chemistry of the lipid, whereas in a mixed bilayer it is set by the surrounding membrane. Apart from the total number of double bonds, which affects the barrier beyond its effect on packing, the tail descriptors and the linkage have small individual effects, presumably because thickness and tail order already carry this information. Area per lipid and experimental main transition temperatures likewise add no resolved information (SI Section S3.5). Lateral packing is captured in the model through thickness and tail order, and local packing around the flip-flopping lipid carries most of the information needed to predict its barrier in a new mixed membrane. Because correlated inputs compensate for one another, the input effects are not additive and should not be read as physical contributions to the barrier.

### 3.5 Potential limitations

Our results come with caveats. The dataset is small for some lipid classes and chemically limited, and the model inputs were chosen with the same cross-validation folds, so the reported errors may be somewhat optimistic for new lipids and membranes. The predictive intervals are also too narrow for PA lipids, so predictions for PA should be treated with caution (Fig. S5). The mixed bilayers contain only ester-linked lipids, so predictions for ether and plasmalogen lipids in mixtures remain untested. The barriers are also specific to Martini 3. They agree qualitatively with native-lipid vesicle experiments, but, assuming equal prefactors, Martini 3 underestimates the differences between lipids and overestimates the effect of cholesterol, and its barriers are 20–70% below CHARMM36m barriers for most lipids. ^25,26^ Capturing the trends with lipid and membrane properties is nevertheless valuable, because experimental barriers are scarce, scattered, and strongly technique-dependent. Systematic atomistic simulations and native-lipid experiments under matched conditions would show which trends, in particular the cholesterol effect, hold beyond Martini 3, and converting barriers into rates would also require the kinetic prefactor.

## 4 Conclusions

We calculated Martini 3 flip-flop free-energy barriers for over 700 membrane–lipid combinations, which, to our knowledge, is the largest such dataset to date. The surrounding membrane affects the barrier as much as the lipid’s own chemistry. Barriers follow the local tail order, increase most strongly with cholesterol, and depend less on the headgroup in mixed membranes. Our Bayesian surrogate model predicts the barriers of withheld lipids and membranes within a few kJ mol^−1^, so an unbiased simulation of a new membrane suffices to estimate the barriers of lipids similar to those in the training data.

Our results have implications for protein-mediated lipid transport. Scramblases, flippases, and floppases act against a composition-dependent background barrier, so their activity and apparent selectivity can be compared across membranes only relative to this barrier, which our dataset and surrogate model can provide. Scramblases that locally deform the surrounding bilayer ^61^ may also lower the barrier outside their hydrophilic groove. Because Martini 3 already overestimates scrambling by membrane proteins, ^25,62^ the biases identified here likely carry over to scramblase simulations.

With the accelerated weight histogram method, a single 15-µs simulation per lipid, without *a priori* knowledge of the system, yielded converged barriers for most lipids. Such highly parallel, biased coarse-grained simulations can thus efficiently generate systematic datasets for training machine learning models that predict how membrane properties change with composition or external conditions. Their success, however, depends critically on the chemical accuracy of the force field.

## Supporting information

Suporting information (SI)

## Author contributions

MV and MJ conceived and designed the study. MV performed all simulations and analyses and wrote the first draft of the manuscript. All authors revised the manuscript and approved the final version.

## Data availability

The simulation files, model files, analysis scripts, and result files underlying this study will be made openly available through the Fairdata IDA service of CSC – IT Center for Science upon submission of the revised manuscript (SI Section S4). The full simulation protocol and model specification are given in SI Sections S1.2 and S3.

## Acknowledgements

We thank CSC–IT Center for Science for computational resources (including LUMI Extreme Scale Access grants “AA-Vesicles” and “PepMemML”). The study was funded by the Research Council of Finland (Academy Research Fellowship grant no. 376628 and PROFI7 funding grant no. 352754).

