## Supplementary material for "Structural Determinants of the Energetic Barrier of Phospholipid Flip-Flop^†^": Suporting information (SI)

**S4 Data files** **12**

#### S1 Simulation details

##### S1.1 Mixed-bilayer compositions

Table S1 lists the numbers of lipids in each leaflet of the 36 mixed bilayers and the flip-flopping lipids of each of them. The initial box height was 12 nm for the single component bilayers and 14 nm for the mixed bilayers, because the box dimensions of the mixed and asymmetric bilayers were expected to fluctuate more (a consequence of the `insane` packing process). The single component bilayers contained 300 lipids per leaflet, and the mixed bilayers were built with lateral dimensions of $14 \times 14$ nm. The rest of the box was filled with Martini water, and in bilayers with a net charge, water beads were replaced with $Na^+$ or $Cl^-$ counterions to neutralize the membrane.

**Table S1** Mixed bilayers. Compositions are the numbers of lipids per leaflet, with one row per leaflet for asymmetric bilayers. The flip-flopping lipids, listed below each composition, are the phospholipid species of the membrane, and those in bold have at least one accepted profile. The PM, ER, Golgi, lysosome, and endosome compositions are adapted from Xu *et al.*,[1] the asymmetric mitochondrial compositions from Brown *et al.*,[2] the *E. coli* composition from Corey *et al.*,[3] and the simplified IMM composition from Makowski *et al.*,[4] retaining their lipid-class proportions with TOCL in place of tetralinoleoyl cardiolipin. Lipids are named as in the Martini 3 lipidome topologies.[5] PM, plasma membrane; IMM and OMM, mitochondrial inner and outer membrane; CHOL, cholesterol; PCER, palmitoyl ceramide; TOCL, tetraoleoyl cardiolipin; DO2B, bis(monoacylglycero)phosphate; BSM, NSM, OSM, PSM, SSM, and XSM, sphingomyelins.

| Membrane | Leaflet | Composition (lipids per leaflet) and flip-flopping lipids | Total |
|---|---|---|---|
| *Phospholipid mixtures* | | | |
| POPC/DOPC 50:50 | | POPC 162, DOPC 162 | 324 |
| | | *Flip-flopping:* **DOPC**, POPC | |

*Table S1 (continued)*

| Membrane | Leaflet | Composition (lipids per leaflet) and flip-flopping lipids | Total |
|---|---|---|---|
| POPC/PLPC 50:50 | | POPC 162, PLPC 162<br>*Flip-flopping:* **PLPC**, **POPC** | 324 |
| POPC/SAPC 50:50 | | POPC 162, SAPC 162<br>*Flip-flopping:* **POPC**, SAPC | 324 |
| POPC/POPE 70:30 | | POPC 226, POPE 97<br>*Flip-flopping:* **POPC**, POPE | 323 |
| POPC/POPS 90:10 | | POPC 291, POPS 32<br>*Flip-flopping:* **POPC**, POPS | 323 |
| POPC/POPS 70:30 | | POPC 226, POPS 97<br>*Flip-flopping:* **POPC**, **POPS** | 323 |
| POPC/POPG 80:20 | | POPC 259, POPG 64<br>*Flip-flopping:* POPC, POPG | 323 |
| POPC/POPA 90:10 | | POPC 291, POPA 32<br>*Flip-flopping:* POPA, POPC | 323 |
| POPC/POPE/POPS 60:30:10 | | POPC 194, POPE 97, POPS 32<br>*Flip-flopping:* **POPC**, POPE, **POPS** | 323 |
| POPC/POPE/POPG 60:30:10 | | POPC 194, POPE 97, POPG 32<br>*Flip-flopping:* **POPC**, POPE, **POPG** | 323 |
| POPC/POPE/PSM 60:30:10 | | POPC 194, POPE 97, PSM 32<br>*Flip-flopping:* **POPC**, POPE, **PSM** | 323 |
| *Cholesterol mixtures* | | | |
| POPC/DOPC/CHOL 50:30:20 | | POPC 162, DOPC 97, CHOL 64<br>*Flip-flopping:* **DOPC**, **POPC** | 323 |
| POPC/PLPC/CHOL 60:20:20 | | POPC 194, PLPC 64, CHOL 64<br>*Flip-flopping:* **PLPC**, **POPC** | 322 |
| POPC/SAPC/CHOL 60:20:20 | | POPC 194, SAPC 64, CHOL 64<br>*Flip-flopping:* **POPC**, **SAPC** | 322 |
| POPC/POPE/CHOL 50:30:20 | | POPC 162, POPE 97, CHOL 64<br>*Flip-flopping:* POPC, POPE | 323 |
| POPC/CHOL 90:10 | | POPC 291, CHOL 32<br>*Flip-flopping:* **POPC** | 323 |
| POPC/CHOL 70:30 | | POPC 226, CHOL 97<br>*Flip-flopping:* **POPC** | 323 |
| DMPC/CHOL 90:10 | | DMPC 291, CHOL 32<br>*Flip-flopping:* DMPC | 323 |
| DMPC/CHOL 80:20 | | DMPC 259, CHOL 64<br>*Flip-flopping:* **DMPC** | 323 |
| DMPC/CHOL 70:30 | | DMPC 226, CHOL 97<br>*Flip-flopping:* **DMPC** | 323 |
| DMPC/CHOL 60:40 | | DMPC 194, CHOL 129<br>*Flip-flopping:* **DMPC** | 323 |
| *Organelle-inspired and bacterial compositions* | | | |
| PM, asymmetric | upper | CHOL 113, POPC 48, PLPC 64, PAPE 9, POPE 32, SSM 32, NSM 12, PCER 9 | 319 |
| | lower | CHOL 93, POPC 22, PLPC 35, PAPE 38, POPE 45, POPI 16, PAPS 35, POPA 9, SSM 9, NSM 16<br>*Flip-flopping:* **NSM**, **PAPE**, **PAPS**, **PLPC**, **POPA**, **POPC**, **POPE**, **POPI**, **SSM** | 318 |
| PM, outer-leaflet control | | CHOL 113, POPC 48, PLPC 64, PAPE 9, POPE 32, SSM 32, NSM 12, PCER 9<br>*Flip-flopping:* **NSM**, **PAPE**, **PLPC**, **POPC**, **POPE**, **SSM** | 319 |
| PM, inner-leaflet control | | CHOL 93, POPC 22, PLPC 35, PAPE 38, POPE 45, POPI 16, PAPS 35, POPA 9, SSM 9, NSM 16<br>*Flip-flopping:* NSM, PAPE, PAPS, PLPC, POPA, POPC, POPE, POPI, SSM | 318 |
| ER | | POPC 51, PLPC 58, SAPC 87, POPE 16, PSPE 25, SAPE 22, SAPI 9, PLPI 9, OLPS 9, PSM 12, POPA 3, CHOL 16<br>*Flip-flopping:* **OLPS**, **PLPC**, PLPI, **POPA**, **POPC**, POPE, PSM, PSPE, **SAPC**, SAPE, SAPI | 317 |
| Golgi | | POPC 51, PLPC 45, SAPC 64, PLPE 12, PSPE 25, SAPE 16, POPI 22, SAPI 6, POPS 12, BSM 38, CHOL 25 | 316 |

*Table S1 (continued)*

| Membrane | Leaflet | Composition (lipids per leaflet) and flip-flopping lipids | Total |
|---|---|---|---|
| | | *Flip-flopping:* BSM, PLPC, PLPE, POPC, POPI, POPS, PSPE, SAPC, SAPE, SAPI | |
| Lysosome | | PLPC 34, SAPC 57, SDPC 19, PLPE 22, SAPE 57, SAPI 19, SDPI 6, SAPS 6, SDPS 3, PSM 9, XSM 9, DO2B 22, CHOL 57 | 320 |
| | | *Flip-flopping:* **DO2B**, **PLPC**, **PLPE**, **PSM**, **SAPC**, SAPE, **SAPI**, **SAPS**, **SDPC**, SDPI, **SDPS**, **XSM** | |
| Endosome | | PLPC 31, SAPC 54, SDPC 19, PLPE 6, PDPE 12, SAPE 22, SAPI 19, SDPI 6, XSM 25, OSM 25, CHOL 101 | 320 |
| | | *Flip-flopping:* OSM, **PDPE**, **PLPC**, **PLPE**, **SAPC**, **SAPE**, **SAPI**, **SDPC**, **SDPI**, **XSM** | |
| IMM, asymmetric | upper | POPC 93, SAPE 116, SAPI 19, POPS 9, TOCL 84 | 321 |
| | lower | POPC 164, SAPE 105, SAPI 14, POPS 8, TOCL 31 | 322 |
| | | *Flip-flopping:* **POPC**, **POPS**, SAPE, SAPI, TOCL | |
| IMM, upper-leaflet control | | POPC 93, SAPE 116, SAPI 19, POPS 9, TOCL 84 | 321 |
| | | *Flip-flopping:* POPC, POPS, SAPE, SAPI, TOCL | |
| IMM, lower-leaflet control | | POPC 164, SAPE 105, SAPI 14, POPS 8, TOCL 31 | 322 |
| | | *Flip-flopping:* POPC, **POPS**, SAPE, SAPI, TOCL | |
| IMM, simplified | | POPC 129, POPE 110, POPI 16, POPS 9, TOCL 58 | 322 |
| | | *Flip-flopping:* **POPC**, POPE, POPI, POPS, TOCL | |
| OMM, asymmetric | upper | POPC 137, SAPE 103, SAPI 16, CHOL 50, PCER 16 | 322 |
| | lower | POPC 168, SAPE 45, SAPI 61, CHOL 48 | 322 |
| | | *Flip-flopping:* **POPC**, **SAPE**, SAPI | |
| OMM, upper-leaflet control | | POPC 137, SAPE 103, SAPI 16, CHOL 50, PCER 16 | 322 |
| | | *Flip-flopping:* **POPC**, **SAPE**, SAPI | |
| OMM, lower-leaflet control | | POPC 168, SAPE 45, SAPI 61, CHOL 48 | 322 |
| | | *Flip-flopping:* POPC, SAPE, **SAPI** | |
| *E. coli* inner membrane | | POPE 217, POPG 74, TOCL 32 | 323 |
| | | *Flip-flopping:* POPE, **POPG**, TOCL | |

#### S1.2 Simulation protocol

Table S2 lists the simulation settings, which were the same for the single component and mixed bilayers. All bilayers were energy-minimized and then pre-equilibrated in six NPT stages with increasing time steps, using a protocol modified from that of CHARMM-GUI Martini Maker.[6] The unbiased and AWH simulations were started independently from the pre-equilibrated bilayers and used the same settings (except the AWH protocol settings block).

The sampled interval of the AWH simulations extended from $-h$ to $+h$ about the centre of mass of the other lipids of the bilayer, with

$$h = \min\left(1.05\, d_\mathrm{f},\ L_z/2 - 0.7\ \mathrm{nm}\right).$$

Here, $d_\mathrm{f}$ is the bilayer thickness in the final structure of the unbiased simulation, measured as the separation of the mean headgroup-marker positions of the two leaflets (the phosphate beads for phospholipids), and $L_z$ is the height of the simulation box. The interval thus spans $2.1\, d_\mathrm{f}$ unless its ends would come closer than 0.7 nm to the periodic limits at $\pm L_z/2$. All flip-flopping lipids of a membrane used the same interval. In asymmetric bilayers, the interval was also symmetric about the reference centre of mass, which can differ from the midpoint between the two leaflets.

#### S1.3 Free-energy profiles and screening

Fig. S1 shows example PMF profiles. The PMF stability screen, described in the main text, combines two criteria, namely the temporal stability of a profile and the agreement between its two side barriers, $B_\mathrm{L}$ and $B_\mathrm{R}$. The symmetry criterion was applied to the single component bilayers, and symmetric composition mixed bilayers. It was not applied to the asymmetric bilayers. For asymmetric membranes, when profiles started from both leaflets of a mixed bilayer were accepted, their barriers were averaged. Table S3 lists the numbers of profiles, membrane–lipid combinations, and host bilayers after each step.

**Table S2** Simulation settings.

| Setting | Value |
|---|---|
| *All stages* | |
| Software and force field | GROMACS 2025.4; Martini 3 with the version 3.2 parameters of the refined lipidome |
| Pair list | Verlet scheme, updated every 20 steps, buffer tolerance 0.005 $\mathrm{kJ\,mol^{-1}\,ps^{-1}}$ per particle |
| Electrostatics | Reaction field, cutoff 1.1 nm, $\varepsilon_\mathrm{r} = 15$, $\varepsilon_\mathrm{rf} = \infty$ |
| Lennard-Jones interactions | Cutoff 1.1 nm, potential shifted to zero at the cutoff |
| *Energy minimization* | |
| Algorithm | Steepest descent in three successive stages of at most 5000 steps, default force tolerance of 10 $\mathrm{kJ\,mol^{-1}\,nm^{-1}}$ |
| First stage | Soft-core nonbonded interactions, cutoffs 2.0 nm |
| *Pre-equilibration and production* | |
| Integrator | Leap-frog |
| Temperature | 310.15 K, stochastic velocity rescaling, coupling time 1 ps, membrane and solvent coupled separately |
| Pressure | 1 bar, semi-isotropic coupling, compressibility $3 \times 10^{-4}$ $\mathrm{bar^{-1}}$ |
| *Pre-equilibration* | |
| Stages 1–5 | 1, 1, 1, 0.75, and 1 ns with time steps of 2, 5, 10, 15, and 20 fs; Berendsen barostat, coupling time 5 ps |
| Stage 6 | 30 ns with a 20 fs time step; Parrinello–Rahman barostat, coupling time 12 ps |
| Velocities | Generated at 310.15 K at the start of stage 1 |
| *Production, unbiased and AWH simulations* | |
| Start | From the pre-equilibrated bilayers, with new velocities generated at 310.15 K |
| Time step | 20 fs |
| Barostat | Stochastic cell rescaling, coupling time 4 ps |
| Unbiased simulations | 10 $\mu$s, coordinates saved every 0.4 ns, descriptors averaged over the final 5 $\mu$s |
| *AWH* | |
| Reaction coordinate | Displacement of the phosphate bead of the flip-flopping lipid along the membrane normal from the centre of mass of the other lipids |
| Bias | Convolved bias, harmonic coupling with $10^4$ $\mathrm{kJ\,mol^{-1}\,nm^{-2}}$, uniform target distribution |
| Settings | Initial error estimate 10 $\mathrm{kJ\,mol^{-1}}$, diffusion estimate $10^{-3}$ $\mathrm{nm^2\,ps^{-1}}$, exponential-to-linear histogram growth |
| Length and output | 15 $\mu$s; pull coordinate saved every 0.8 ns, coordinates and bias every 8 ns |

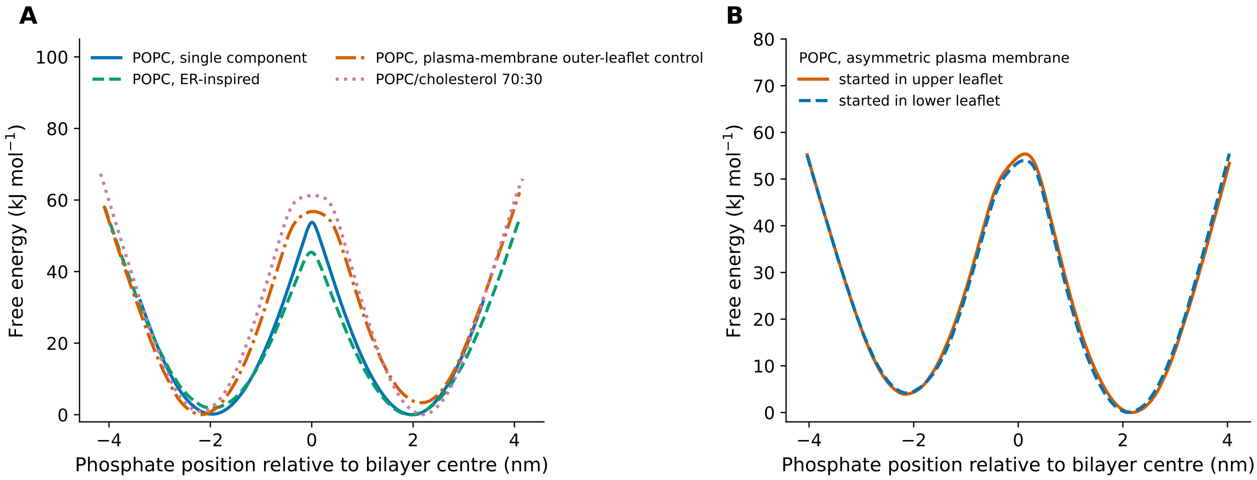

**Fig. S1** Example free-energy profiles (PMFs). (A) POPC in its single component bilayer, the ER composition, the outer-leaflet PM control, and POPC/CHOL 70:30. (B) POPC in the asymmetric PM composition, started from either leaflet, with the upper (outer) leaflet at positive positions. The two basins in (B) differ by about 4 $\mathrm{kJ\,mol^{-1}}$, which is the leaflet preference that $B_\mathrm{ave}$ averages out. The minimum of each profile is set to zero.

**Table S3** Screening of the free-energy profiles. The PMF stability screen combines temporal stability and side-barrier symmetry, and profiles that did not converge count as not temporally stable. Each single component bilayer has one profile. In mixed bilayers, a membrane–lipid combination can have profiles started from both leaflets, and a host bilayer contains several combinations. The 37 of the 645 simulated single component bilayers without complete structural data are not counted, and seven gel-like bilayers passed the PMF stability screen. The model dataset excludes the eight single component PI, SM, and BMP bilayers (DO2B, DO3B, MSM, OSM, PGSM, PLPI, POPI, and SAPI) and the 14 mixed-bilayer combinations of DO2B, NSM, POPI, PSM, SAPI, SDPI, SSM, and XSM.

| | Single component bilayers | Mixed profiles | Mixed combinations | Host bilayers |
|---|---|---|---|---|
| Simulated profiles | 608 | 152 | 138 | 36 |
| Converged (temporally stable) | 473 | 82 | 76 | 30 |
| Passed the PMF stability screen | 436 | 80 | 74 | 29 |
| Analysed dataset (gel-like bilayers removed) | 429 | 80 | 74 | 29 |
| Model dataset (PA, PC, PE, PG, and PS lipids) | 421 | 64 | 60 | 28 |

Gel-like bilayers were identified among the single component bilayers only. The area per lipid, bilayer thickness, and bulk and interfacial tail order of the 608 single component bilayers with complete structural data were standardized, and their first principal component, which explains 89.25% of the variance, served as a packing coordinate. A two-component Gaussian mixture was fitted to this coordinate with 50 initializations and a fixed random seed. The 123 bilayers with a probability of at least 0.5 for the ordered component were classified as gel-like and the remaining 485 as fluid-like (Fig. S2). The gel-like bilayers were excluded from all analyses. The mixed bilayers were projected onto the same coordinate but not excluded, because high tail order in cholesterol-containing membranes does not indicate a gel phase. Notably, seven combinations in four cholesterol-rich mixed bilayers fall into the gel-like region.

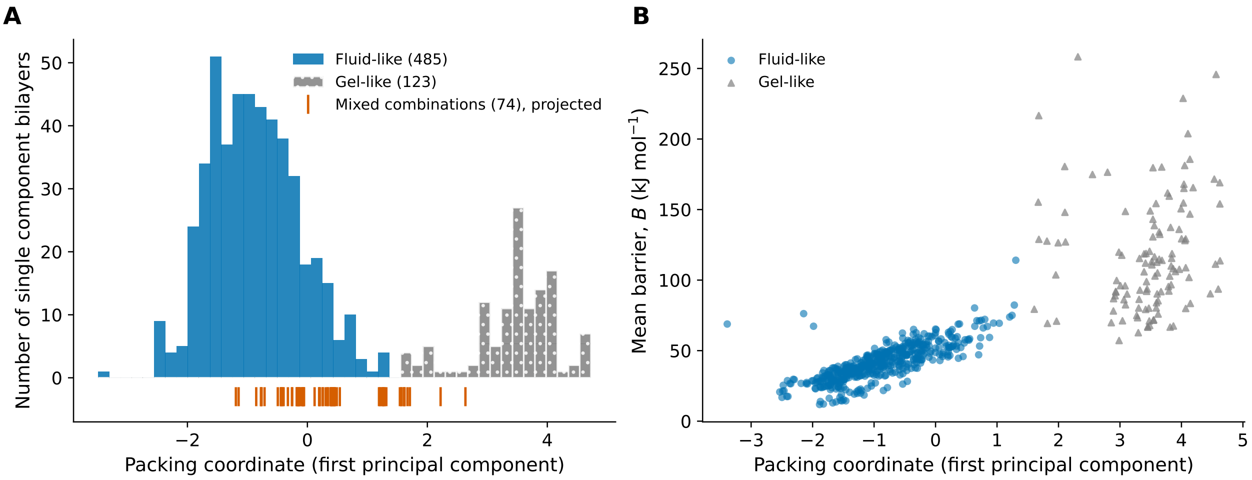

**Fig. S2** Identification of gel-like single component bilayers. (A) Packing coordinate of the 608 single component bilayers, classified as fluid-like or gel-like, with the 74 accepted mixed-bilayer combinations projected onto the same coordinate for comparison (ticks). (B) Mean barrier of the single component bilayers against the packing coordinate.

The screening does not remove profiles at random, which qualifies some conclusions. The fluid-like single component bilayers excluded by the PMF stability screen have a lower mean barrier than the retained ones (37 against 42 kJ mol$^{-1}$), so the screen slightly raises the mean barrier of the dataset. In the mixed bilayers, PE profiles passed the screen less often than PC profiles (11 of 34 against 41 of 56). The model dataset contains 89 PA, 86 PC, 73 PE, 86 PG, and 87 PS single component bilayers (142 ester, 140 ether, and 139 plasmalogen), and its 60 mixed-bilayer combinations comprise 18 ester-linked lipids in nine tail families.

#### S2 Additional results

##### S2.1 Structural descriptors and barrier shifts

The structural descriptors are strongly correlated with one another. In single component bilayers, bulk and interfacial tail order correlate with $r = 0.91$, and area per lipid and bulk tail order with $r = -0.91$. In mixed bilayers, the two order parameters correlate with $r = 0.98$. Fig. 3 of the main text shows how the barrier correlates with each descriptor.

The barrier shift of a lipid moved from its single component bilayer into a mixed bilayer follows the accompanying change in bulk tail order ($r = 0.83$, 95% CI 0.77–0.90) and, more weakly, the changes in interfacial tail order ($r = 0.77$), area per lipid ($r = -0.65$), and thickness ($r = 0.62$; Fig. S3). The correlation with bulk tail order is unchanged when the five shifts in the gel-like region are removed ($r = 0.83$, 95% CI 0.76–0.91).

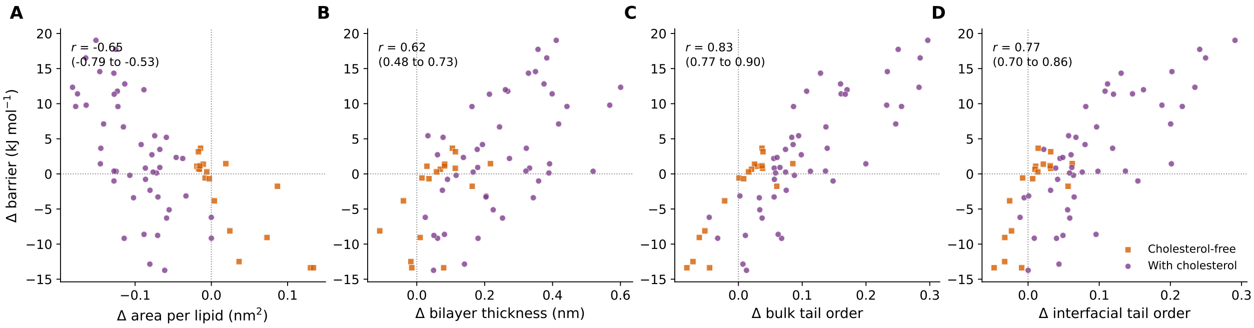

**Fig. S3** Barrier shift on moving a lipid from its single component bilayer into a mixed bilayer against the accompanying change in (A) area per lipid, (B) bilayer thickness, (C) bulk tail order, and (D) interfacial tail order, for 65 membrane–lipid combinations. Squares show cholesterol-free and circles cholesterol-containing mixed bilayers. Area changes compare the box area with the local Voronoi area and are therefore approximate.

#### S2.2 Chemical and membrane comparisons

Table S4 lists the numbers of pairs, medians, and intervals of the matched comparisons in single component bilayers that are discussed in the main text. Lipids with 18:1(11-*cis*) chains are omitted from Fig. 4B for clarity but are included in this table and in all other analyses.

Table S5 lists the 13 comparisons in which a lipid and its PC analogue share a mixed bilayer, together with the barrier difference between their single component bilayers. Within a mixed bilayer, the two lipids sample practically the same local environment, as their local tail order parameters differ by less than 0.003 and their local thicknesses by less than 0.005 nm. These comparisons thus isolate the effect of the headgroup. Between single component bilayers, in contrast, the two lipids also form different membranes. For example, the bulk tail order is 0.54 in the POPE bilayer but 0.40 in the POPC bilayer, so their barrier difference of 17.6 $\mathrm{kJ\,mol^{-1}}$ also includes the effect of the more ordered POPE membrane. The local environment also differs strongly between mixed bilayers, so the same headgroup contrast can vary between them. The bulk tail order around POPC, for example, is 0.55–0.60 in the PM compositions but 0.37 in the ER composition.

**Table S4** Matched chemical comparisons in single component bilayers. Each comparison changes one feature and keeps the others fixed. Groups are the distinct tail compositions, including double-bond positions, and the 95% intervals are from cluster bootstrapping over these groups. For the tail comparisons, the range of differences is given instead.

| Comparison | Pairs | Groups | Median ($\mathrm{kJ\,mol^{-1}}$) | 95% CI or range | Positive differences |
|---|---|---|---|---|---|
| PE–PC | 65 | 28 | 12.69 | 10.95 to 13.93 | 65 of 65 |
| PS–PC | 78 | 29 | 7.19 | 6.55 to 7.62 | 78 of 78 |
| PG–PC | 76 | 29 | 2.05 | 0.56 to 2.84 | 60 of 76 |
| PA–PC | 80 | 30 | −4.88 | −5.78 to −4.31 | 3 of 80 |
| Ether–ester | 129 | 31 | −1.88 | −2.14 to −1.52 | 2 of 129 |
| Plasmalogen–ester | 127 | 31 | 0.29 | 0.19 to 0.43 | 94 of 127 |
| More double bonds, same carbon counts | 541 | | −11.11 | range −43.09 to 3.79 | 11 of 541 |
| Two more carbons per chain, symmetric monounsaturated | 78 | | 3.88 | range −0.83 to 15.37 | 74 of 78 |
| 18:1 double bond at position 11 instead of 9 | 15 | | −4.29 | range −6.63 to −1.70 | 0 of 15 |

**Table S5** Headgroup effect relative to PC within the same mixed bilayer and between the single component bilayers of the same two lipids.

| Mixed bilayer | Lipid | PC analogue | Within the mixed bilayer ($\mathrm{kJ\,mol^{-1}}$) | Between single component bilayers ($\mathrm{kJ\,mol^{-1}}$) |
|---|---|---|---|---|
| Endosome | PLPE | PLPC | +12.58 | +12.13 |
| Lysosome | PLPE | PLPC | +8.85 | +12.13 |
| ER | POPA | POPC | −4.07 | −7.04 |
| PM, asymmetric | POPA | POPC | −5.63 | −7.04 |
| PM, asymmetric | POPE | POPC | +9.82 | +17.58 |
| PM, outer-leaflet control | POPE | POPC | +7.52 | +17.58 |
| POPC/POPE/POPG 60:30:10 | POPG | POPC | −0.08 | +0.05 |
| IMM, asymmetric | POPS | POPC | +6.55 | +6.54 |
| POPC/POPE/POPS 60:30:10 | POPS | POPC | +2.79 | +6.54 |
| POPC/POPS 70:30 | POPS | POPC | +2.41 | +6.54 |
| Endosome | SAPE | SAPC | +9.28 | +13.14 |
| Lysosome | SAPS | SAPC | +2.97 | +7.93 |
| Lysosome | SDPS | SDPC | +5.36 | +7.55 |

Fig. S4 shows the barriers of the five lipids accepted in at least three organelle-inspired compositions. The barriers of POPC and PLPC are highest in the PM compositions, which have the highest local bulk tail order, whereas those of PLPC and SAPC change little between the ER, lysosome, and endosome compositions. Lipids that share a membrane thus follow similar trends. Membranes with similar local tail order can still differ. SAPI, for example, has barriers of 53.3 and 60.1 $\mathrm{kJ\,mol^{-1}}$ in the endosome composition and the lower-leaflet OMM control, although its local bulk and interfacial tail order are 0.42 and 0.51 in the endosome composition and 0.43 and 0.50 in the OMM control.

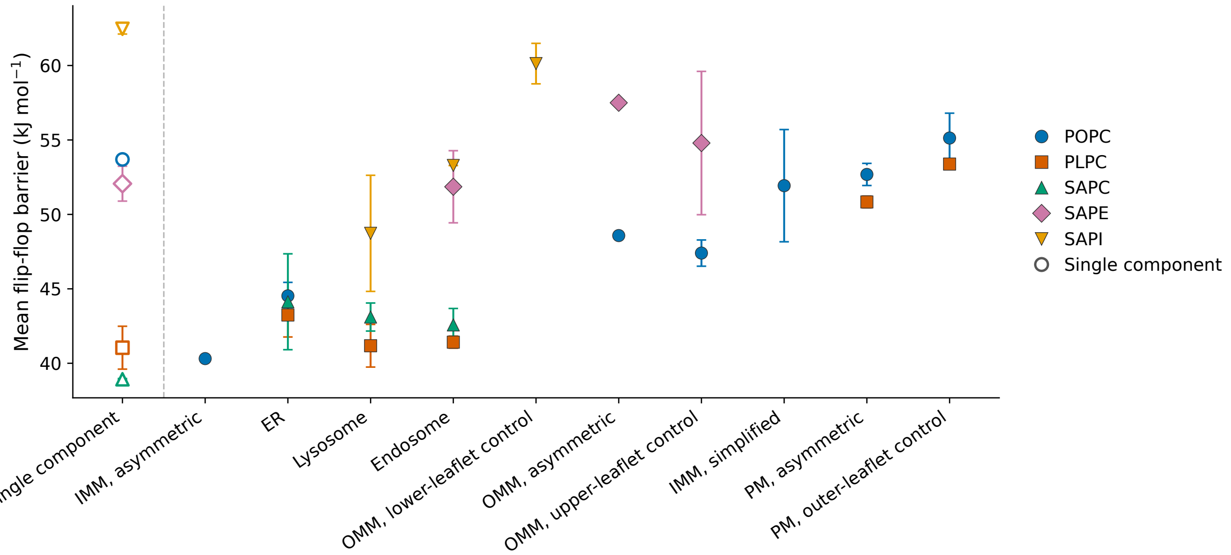

**Fig. S4** Barriers of the lipids accepted in at least three organelle-inspired compositions, in their single component bilayers (open symbols) and in each composition (filled symbols). The compositions are ordered by the mean bulk tail order around their accepted lipids, from 0.36 in the asymmetric IMM composition to 0.60 in the outer-leaflet PM control. Asymmetric compositions give the direction-averaged barrier $B_{\mathrm{ave}}$. The error bars span $B_{\mathrm{L}}$ and $B_{\mathrm{R}}$ of the profile in symmetric bilayers and the barriers of the two profiles started from opposite leaflets in asymmetric bilayers.

##### S2.3 Asymmetric bilayers

In an asymmetric bilayer, the two side barriers start from basins in different leaflet environments. The left and right basins, at negative and positive $z$, lie in the lower and upper leaflets, respectively, and in the PM the upper leaflet is the outer leaflet. $B_{\mathrm{L}}$ and $B_{\mathrm{R}}$ are thus the directional barriers for flip-flop from the lower and the upper leaflet. Their difference, $B_{\mathrm{L}} - B_{\mathrm{R}} = \Delta G(z_{\mathrm{R}}) - \Delta G(z_{\mathrm{L}})$, is the leaflet preference of the lipid, which $B_{\mathrm{ave}}$ averages out. Table S6 lists $B_{\mathrm{L}}$ and $B_{\mathrm{R}}$ of the 13 accepted combinations. When profiles were started from both leaflets, their side barriers were averaged, and the two profiles agree within 3.1 $\mathrm{kJ\,mol^{-1}}$, except for $B_R$ of SSM (5.9 kJ mol$^{-1}$). The two leaflets of each asymmetric bilayer contain almost equal numbers of lipids although their compositions differ (Table S1).

The directional barriers agree closely with the available leaflet-matched symmetric controls. In all nine comparisons, $B_L$ or $B_R$ lies within 2.4 kJ mol$^{-1}$ of the barrier in the lower- or upper-leaflet control, respectively. The direction-averaged barrier $B_{ave}$ agrees less closely and lies within 3 kJ mol$^{-1}$ of the controls in eight of the nine comparisons.

Excluding the asymmetric bilayers from training has little effect on the model. Because $B_{ave}$ of an asymmetric bilayer averages two different directions, we refitted the model on the same folds without the 10 asymmetric combinations of the model dataset. The model then predicts the other 50 mixed-bilayer combinations with an RMSE of 3.8 kJ mol$^{-1}$, compared with 3.6 kJ mol$^{-1}$ when the asymmetric combinations are included in training (difference +0.20 kJ mol$^{-1}$, 95% CI −0.06 to +0.37), and the family-test RMSE changes by −0.03 kJ mol$^{-1}$ (95% CI −0.08 to +0.01). The asymmetric combinations themselves are predicted with an RMSE of 4.9 kJ mol$^{-1}$ in the membrane test.

**Table S6** Directional barriers in the asymmetric bilayers (kJ mol$^{-1}$). $B_L$ and $B_R$ are the barriers for flip-flop from the lower and upper leaflet, and their difference is the leaflet preference. The controls are the barriers in the leaflet-symmetric bilayers with the composition of the starting leaflet. $B_{ave}$ is the direction-averaged barrier used in the main text.

| Asymmetric bilayer | Lipid | $B_L$ | Lower-leaflet control | $B_R$ | Upper-leaflet control | $B_L - B_R$ | $B_{ave}$ |
|---|---|---|---|---|---|---|---|
| IMM, asymmetric | POPC | 50.62 | | 30.01 | | +20.62 | 40.31 |
| IMM, asymmetric | POPS | 50.45 | 51.17 | 43.28 | | +7.17 | 46.86 |
| OMM, asymmetric | POPC | 48.00 | | 49.15 | 47.40 | −1.15 | 48.58 |
| OMM, asymmetric | SAPE | 62.56 | | 52.43 | 54.79 | +10.13 | 57.50 |
| PM, asymmetric | NSM | 59.39 | | 62.29 | 63.29 | −2.90 | 60.84 |
| PM, asymmetric | PAPE | 60.94 | | 61.17 | 63.00 | −0.23 | 61.05 |
| PM, asymmetric | PAPS | 49.38 | | 56.11 | | −6.73 | 52.74 |
| PM, asymmetric | PLPC | 49.42 | | 52.25 | 53.38 | −2.83 | 50.84 |
| PM, asymmetric | POPA | 45.49 | | 48.62 | | −3.13 | 47.05 |
| PM, asymmetric | POPC | 50.66 | | 54.71 | 55.13 | −4.04 | 52.68 |
| PM, asymmetric | POPE | 63.70 | | 61.30 | 62.65 | +2.40 | 62.50 |
| PM, asymmetric | POPI | 64.14 | | 70.66 | | −6.52 | 67.40 |
| PM, asymmetric | SSM | 62.52 | | 60.53 | 62.73 | +1.98 | 61.52 |

#### S2.4 Comparison with experiments

Table S7 compares the barrier differences implied by native-lipid vesicle experiments with those calculated here. Activation free energies derived from measured rates with transition-state theory, such as 104 kJ mol$^{-1}$ for DMPC at 37 °C,[7] are not directly comparable with PMF barriers, because they assume the prefactor $k_B T/h$ of transition-state theory, where $h$ is the Planck constant. The prefactor of flip-flop is likely much smaller, and each ten-fold decrease in the prefactor lowers the implied barrier by 5.9 kJ mol$^{-1}$ at 310.15 K.

**Table S7** Barrier differences $B_1 - B_2$ between the first and the second system of each comparison, implied by flip-flop half-times in native-lipid vesicles (100 nm extruded vesicles, 150 mM NaCl, 37 °C)[8] and calculated here (kJ mol$^{-1}$). The implied differences assume equal prefactors, $B_1 - B_2 = RT \ln(t_{1/2,1}/t_{1/2,2})$, so a negative value means that the first system flips faster.

| Comparison | Measured half-times | Implied $B_1 - B_2$ | Martini 3, this work |
|---|---|---|---|
| POPA – POPC | ∼7 h and >1000 h | $< -12.8$ | −7.0 |
| DMPC – POPC | 350 min and >1000 h | $< -13.3$ | −6.3 |
| DMPC with 20 mol% cholesterol – DMPC | 1300 min and 350 min | +3.4 | +12.0 |
| DMPC with 40 mol% cholesterol – DMPC | >7000 min and 350 min | $> +7.7$ | +19.0 |

### S3 Bayesian model

#### S3.1 Overview

The surrogate model predicts the barrier of a lipid from its chemistry and from the packing of the membrane around it. The chemistry enters through the headgroup and linkage and through five numerical descriptors of the tails, and the packing through the thickness and the bulk and interfacial tail order from the unbiased simulations. Most relationships are linear, but smooth splines allow nonlinearity in the tail length, thickness, and interfacial order responses. Each tail family and each mixed bilayer has its own offset, so that the shared behaviour of related observations is not attributed to the descriptors. No coefficient belongs to an individual lipid. Mixed bilayers also have their own corrections to the structural slopes and their own residual scatter. The priors keep the corrections, that is, the interactions, the spline terms, and the mixed-bilayer slopes, close to zero unless the data clearly support them. The model describes the logarithm of the barrier, so its inputs change the barrier by a factor rather than by a fixed amount.

#### S3.2 Model and priors

The model of eqn (2) of the main text is a linear regression for $y_i = \ln(B_i/50)$, where $B_i$ is the mean barrier $B_{\mathrm{ave}}$ of observation $i$ in kJ mol$^{-1}$, averaged over the accepted profiles of a mixed-bilayer combination. The division by 50 kJ mol$^{-1}$ only sets the reference point of the logarithm. With $m_i = 1$ for a mixed-bilayer combination and $m_i = 0$ for a single component bilayer, observation $i$ is assumed to follow

$$y_i \sim \mathcal{N}\left(\eta_i,\ \sigma^2 r^{2m_i}\right), \qquad r \in \{1, 2, 4, 8\}. \tag{S1}$$

The residual standard deviation is thus $\sigma$ for single component bilayers and $r\sigma$ for mixed bilayers. The ratio $r$ is not known in advance, so four values with equal prior weights are considered. The mean is

$$\eta_i = \alpha + \boldsymbol{\beta}_{\mathrm{c}}^{\mathsf{T}}\mathbf{c}_i + \boldsymbol{\beta}_{\mathrm{s}}^{\mathsf{T}}\mathbf{s}_i + h_{H_i} + \ell_{L_i} + \gamma_{H_i} s_{i,\mathrm{I}} + \omega\, c_{i,\mathrm{C}}\, s_{i,d} + \boldsymbol{\lambda}^{\mathsf{T}}\mathbf{p}_i + u_{f_i} + m_i\left(\delta + \boldsymbol{\kappa}^{\mathsf{T}}\mathbf{s}_i + v_{g_i}\right), \tag{S2}$$

where Table S8 explains each term. Here, $c_{i,\mathrm{C}}$ is the total number of tail carbons, $s_{i,d}$ the thickness, $s_{i,\mathrm{I}}$ the interfacial tail order, and $\mathbf{p}_i$ contains the five spline functions. The total numbers of tail carbons and double bonds are summed over both chains, and the double-bond asymmetry is the absolute difference in the number of double bonds between the two chains. The unsaturation position is the mean position of the unsaturated tail beads along their chains, from 0 at the first tail bead to 1 at the last, and is 0 for saturated lipids. The mapped tail bending is the mean of $(180^\circ - \theta_0)/180^\circ$ over the equilibrium angles $\theta_0$ of the tail angle potentials in the Martini topology. All numerical descriptors are standardized to zero mean and unit standard deviation over the training data.

The splines allow the barrier to curve with total tail carbons, thickness, and interfacial tail order, whereas the linear terms describe the overall trends. Each spline is a natural cubic spline with knots at the 10th, 35th, 65th, and 90th percentiles of the training data, and it is linear beyond the outer knots. Each spline function is made uncorrelated with the linear term of its descriptor and scaled to unit standard deviation, so that it describes only the curvature. Total tail carbons take few distinct values, so two of their knots coincide and leave one spline function, whereas thickness and interfacial tail order have two spline functions each. The knots and splines, like the standardization, were determined from the training data of each cross-validation fold only.

**Table S8** Terms of the mean $\eta_i$ in eqn (S2). The indices $H_i$, $L_i$, $f_i$, and $g_i$ denote the headgroup, linkage, tail family, and mixed bilayer of observation $i$, and $m_i = 1$ for mixed-bilayer combinations and 0 otherwise.

| Term | Physical definition | Implementation in the model |
|---|---|---|
| Intercept, $\alpha$ | Typical barrier of a single component bilayer | Constant |
| Chemical descriptors, $\mathbf{c}_i$ | Total tail carbons, total double bonds, double-bond asymmetry between the chains, unsaturation position, and mapped tail bending | Linear slopes $\boldsymbol{\beta}_\mathrm{c}$ |
| Structural descriptors, $\mathbf{s}_i$ | Thickness, bulk tail order, and interfacial tail order of the surrounding membrane | Linear slopes $\boldsymbol{\beta}_\mathrm{s}$ |
| Headgroup, $h_H$ | PA, PC, PE, PG, or PS | One offset per headgroup |
| Linkage, $\ell_L$ | Ester, ether, or plasmalogen | One offset per linkage |
| Headgroup–order correction, $\gamma_H$ | Headgroups can follow interfacial tail order differently | Correction to the slope of interfacial tail order for each headgroup |
| Carbon–thickness interaction, $\omega$ | The effect of tail length can depend on the thickness | Product of total tail carbons and thickness |
| Curvature, $\boldsymbol{\lambda}$ | Smooth curves that allow the relationships to bend | Five spline functions, one in total tail carbons and two each in thickness and interfacial tail order |
| Tail family, $u_f$ | Shared deviation of all lipids with the same tails | One offset per tail family |
| Mixed-bilayer offset, $\delta$ | Systematic difference between mixed and single component bilayers | Common offset for all mixed-bilayer combinations |
| Mixed-bilayer slopes, $\boldsymbol{\kappa}$ | Structure can act differently in mixed bilayers | Corrections to the three structural slopes |
| Mixed bilayer, $v_g$ | Shared deviation of all lipids in one mixed bilayer | One offset per mixed bilayer |

All coefficients are collected in the vector $\boldsymbol{\phi}$. The priors are

$$p(\sigma^2) \propto \left(\sigma^2\right)^{-(a_0+1)} \exp\left(-\frac{b_0}{\sigma^2}\right), \qquad \boldsymbol{\phi} \mid \sigma^2 \sim \mathcal{N}\left(\mathbf{0},\ \sigma^2\mathbf{D}\right), \qquad D_{jj} = \frac{\tau_j^2}{0.01}, \tag{S3}$$

with $a_0 = 2$ and $b_0 = 0.01$. The prior mean of $\sigma^2$ is $b_0/(a_0 - 1) = 0.01$, which corresponds to a residual standard deviation of about 10% of the barrier, and at this value the prior standard deviation of coefficient $\phi_j$ is $\tau_j$ (Table S9). These zero-centred priors act like the penalty of ridge regression. Coefficients with small prior scales are pulled more strongly towards zero, so that the corrections stay small unless the data require them. The prior scales are fixed rather than estimated from the data. The headgroup and linkage offsets use one indicator per category alongside the intercept and are thus separated only through their priors.

**Table S9** Prior scales $\tau_j$ of the coefficients, in units of the log barrier per standardized input.

| Coefficients | $\tau_j$ |
|---|---|
| $\alpha$ | 0.5 |
| $\boldsymbol{\beta}_\mathrm{s}$, $h_H$ | 0.3 |
| $\boldsymbol{\beta}_\mathrm{c}$, $\delta$ | 0.2 |
| $v_g$ | 0.15 |
| $\ell_L$, $u_f$ | 0.1 |
| $\gamma_H$, $\omega$, $\boldsymbol{\lambda}$, $\boldsymbol{\kappa}$ | 0.08 |

#### S3.3 Predictions and their uncertainty

Because the normal–inverse-gamma priors are conjugate, the fitted coefficients and the predictions are calculated directly, without sampling. Each value of $r$ is weighted by its marginal likelihood, which gives weights of 0.03 and 0.97 to $r = 1$ and $r = 2$ for the final model and negligible weights to $r = 4$ and 8. The prediction for a new observation is a weighted mixture of Student $t$ distributions. The predicted barrier is its median, and the 95% predictive interval covers four sources of uncertainty. These are the remaining scatter of the barriers around the model, the uncertainty of the fitted coefficients, the unknown offset of a tail family or mixed bilayer that is absent from the training data, and the block standard errors of the structural descriptors of the new observation, propagated to first order. The model uses no separate error for each PMF, so the scatter term also absorbs the sampling error of the profiles. The error of the force field itself is not included.

##### S3.4 Validation

Predictive accuracy was assessed with the family and membrane tests of the main text.[9] The family test also predicts the mixed-bilayer combinations of the withheld tail families, and the mixed bilayers that contain these lipids are then removed from training together with their leaflet-symmetric controls. Accuracy is the RMSE of the predicted median, with 95% intervals from 3000 bootstrap resamples of the withheld groups. Calibration is the coverage, that is, the fraction of calculated barriers that fall inside the predictive interval. Differences between models are from 3000 paired group-bootstrap resamples on the same folds.

Table S10 summarizes the held-out performance. The training RMSEs are 1.20 and 2.55 $\mathrm{kJ\,mol^{-1}}$ for single component and mixed bilayers. The predictive intervals are slightly too narrow for single component bilayers and slightly too wide for mixed bilayers (Fig. S5A). The PA barriers are predicted about as accurately as those of the other headgroups (RMSE 2.78 against 2.47–3.19 $\mathrm{kJ\,mol^{-1}}$), but their intervals are the narrowest, and only 80.9% of the PA barriers lie inside their 95% intervals (Fig. S5B). The model is therefore overconfident for PA. Lipids with two chains of equal length, which span a wider range of total tail carbons (16–44 against 30–40), are predicted less accurately than lipids with unequal chains (RMSE 3.4 against 1.8 $\mathrm{kJ\,mol^{-1}}$).

**Table S10** Held-out performance of the final model. Coverage is the percentage of calculated barriers inside the 50, 80, and 95% predictive intervals.

| Test | $n$ (groups) | RMSE ($\mathrm{kJ\,mol^{-1}}$; 95% CI) | $R^2$ | Coverage at 50/80/95% (%) | Mean 95% width ($\mathrm{kJ\,mol^{-1}}$) |
|---|---|---|---|---|---|
| Family test, single component bilayers | 421 (35) | 2.81 (2.19–3.40) | 0.94 | 47.5 / 74.8 / 91.7 | 9.4 |
| Membrane test, mixed-bilayer combinations | 60 (25) | 3.81 (2.83–4.41) | 0.67 | 61.7 / 88.3 / 96.7 | 19.4 |
| Family test, mixed-bilayer combinations | 60 (9) | 4.76 | 0.49 | 56.7 / 91.7 / 100.0 | 27.4 |

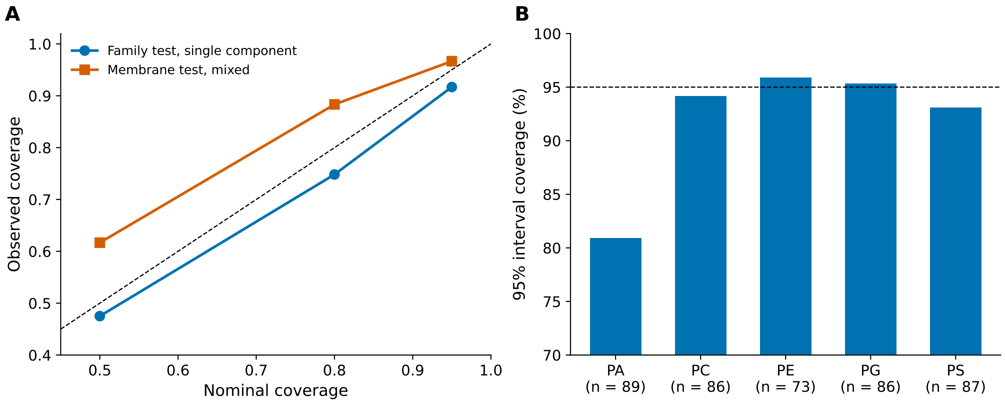

**Fig. S5** Calibration of the final model. (A) Observed against nominal coverage of the 50, 80, and 95% predictive intervals in the family test (single component bilayers) and the membrane test (mixed bilayers). (B) Coverage of the 95% intervals by headgroup in the family test. Dashed lines mark equality in (A) and 95% in (B).

Mixed bilayers in training are essential for predicting new membranes (Table S11). The model trained only on single component bilayers covers every mixed observation, but only because its intervals are very wide (mean width 45.9 $\mathrm{kJ\,mol^{-1}}$). Using the single component barrier of the same lipid gives about twice the error of the final model.

**Table S11** Predictions of the 60 mixed-bilayer combinations with and without mixed bilayers in training. Differences are paired group-bootstrap intervals over the 25 mixed-bilayer groups.

| Prediction | RMSE ($kJ\,mol^{-1}$) | Difference from the final model ($kJ\,mol^{-1}$; 95% CI) |
|---|---|---|
| Final model (membrane test) | 3.81 | — |
| Final model trained on single component bilayers only | 5.94 | +2.13 (+1.33 to +2.91) |
| Single component barrier of the same lipid | 7.95 | +4.14 (+2.97 to +5.51) |

##### S3.5 Input removal and excluded inputs

To measure what each input adds, we removed it, fitted the model again on the same folds, and recorded how much the predictions worsened. Each input was removed together with its spline, interaction, and mixed-slope terms. The tail-family offsets were kept, except when all chemical inputs were removed. The inputs were selected on the same cross-validation folds. An earlier version of the model also included chain-length asymmetry, which was removed because its removal lowered the RMSE in both tests (by 0.10 and 0.04 $kJ\,mol^{-1}$ in the family and membrane tests). Fig. 5C of the main text shows the change for each input. Removing the lipid linkage information changes the RMSE by $-0.07$ $kJ\,mol^{-1}$ (95% CI $-0.37$ to $+0.16$) in the family test and by $+0.05$ $kJ\,mol^{-1}$ ($-0.09$ to $+0.19$) in the membrane test. The linkage was nevertheless retained, so that ester, ether, and plasmalogen lipids can differ in their barriers, as they do at matched headgroup and tails (Table S4).

The effects of additional inputs were also tested. Area per lipid is not an input of the model. Adding it as a fourth structural descriptor, with the same prior scale as the other structural descriptors, changes the membrane-test RMSE by $+0.17$ $kJ\,mol^{-1}$ (95% CI $-0.51$ to $+0.73$). Experimental main transition temperatures are available for 28 ester lipids of the panel.[10–12] They correlate with the barrier ($r = 0.65$) and with interfacial tail order ($r = 0.88$). In a ridge regression on the headgroup and the four structural descriptors of these lipids, validated by withholding one tail family at a time, adding the transition temperature lowers the RMSE by only 0.07 $kJ\,mol^{-1}$ (95% CI $-0.12$ to $+0.18$).

#### S4 Data files

The model code, analysis scripts, result files, and simulation input files will be made openly available through the Fairdata IDA service of CSC – IT Center for Science upon submission of the revised manuscript. The model code performs the fits, cross-validation tests, and predictions reported here. The result files contain the barriers, screening outcomes, and structural descriptors of simulated membranes, together with the free-energy profiles and the model predictions.
